# Neuropeptide Modulation of Bidirectional Internetwork Synapses

**DOI:** 10.1101/2023.12.15.571956

**Authors:** Savanna-Rae H Fahoum, Dawn M Blitz

## Abstract

Oscillatory networks underlying rhythmic motor behaviors, and sensory and complex neural processing, are flexible, even in their neuronal composition. Neuromodulatory inputs enable neurons to switch participation between networks, or participate in multiple networks simultaneously. Neuromodulation of internetwork synapses can both recruit and coordinate a switching neuron in a second network. We previously identified an example in which a neuron is recruited into dual-network activity via peptidergic modulation of intrinsic properties. We now ask whether the same neuropeptide also modulates internetwork synapses for internetwork coordination. The crab (*Cancer borealis*) stomatogastric nervous system contains two well-defined feeding-related networks (pyloric, food filtering, ∼1 Hz; gastric mill, food chewing, ∼0.1 Hz). The projection neuron MCN5 uses the neuropeptide Gly^1^-SIFamide to recruit the pyloric-only LPG neuron into dual pyloric plus gastric mill-timed bursting via modulation of LPG’s intrinsic properties. Descending input is not required for a coordinated rhythm, thus intra-network synapses between LPG and its second network must underlie coordination among these neurons. However, synapses between LPG and gastric mill neurons have not been documented. Using two-electrode voltage clamp recordings, we found that graded synaptic currents between LPG and gastric mill neurons (LG, IC, DG) were primarily negligible in saline, but were enhanced by Gly^1^-SIFamide. Further, LPG and gastric mill neurons entrain each other during Gly^1^-SIFamide application, indicating bidirectional, functional connectivity. Thus, a neuropeptide mediates neuronal switching through parallel actions, modulating intrinsic properties in a switching neuron to recruit it into a second network and as shown here, also modulating bidirectional internetwork synapses for coordination.

**New and Noteworthy:** Neuromodulation can enable neurons to be simultaneously coordinated with separate networks. Both recruitment into, and coordination with, a second network can occur via modulation of internetwork synapses. Alternatively, recruitment can occur via modulation of intrinsic ionic currents. We find that the same neuropeptide previously determined to modulate intrinsic currents also modulates bidirectional internetwork synapses that are typically ineffective. Thus, complementary modulatory peptide actions enable recruitment and coordination of a neuron into a second network.

## Introduction

Central pattern generators (CPGs) are flexible neuronal networks that each produce different versions of particular rhythmic motor behaviors such as walking, breathing, and chewing (1–5). CPG flexibility includes neuronal switching, in which a neuron changes participation from its “home” network into another, or multiple networks simultaneously, via neuromodulation of synaptic and/or intrinsic properties (6–12). Neuronal switching occurs in invertebrate and vertebrate CPG motor networks, other local oscillatory networks such as in the hippocampus, and is proposed to contribute to larger-scale changes in functional connectivity (10, 13–18). Neuromodulator inputs such as cholinergic basal forebrain and locus coeruleus neurons are implicated in regulating network dynamics on multiple temporal and spatial scales (14, 15, 17, 19). To fully understand how neuromodulatory inputs alter neuronal participation, it is necessary to identify the full complement of modulatory actions enabling a switching neuron to generate activity at the same frequency as a new network and with the correct timing relative to other neurons.

Modulation of internetwork synapses onto a switching neuron recruits a neuron to generate activity at the correct frequency and also enables the correct relative timing (6, 8, 20). In such instances, the switching neuron typically follows the pattern imposed on it, but does not contribute to generating the output pattern in its new network. More recent examples indicate that modulation of intrinsic properties can also regulate neuronal switching (10–12). We found that a neuron recruited to generate activity at a second network frequency via modulation of intrinsic properties can coordinate activity of neurons in the network into which it is recruited (21). For a switching neuron to be able to coordinate activity in the new network, one possibility is that there are functional synapses already present, and bursting of the switching neuron at the second frequency simply has effects via those existing synapses. An alternative possibility is that there are not functional synapses to mediate coordination under unmodulated conditions, and modulation of synapses is required to enable coordination between a switching neuron and other neurons in its new network. This suggests that in contrast to modulation of internetwork synapses enabling a neuron to generate activity at the correct frequency and with the correct relative timing, modulation of intrinsic properties may require complementary modulatory actions. Here, we tested the hypothesis that modulation of synapses occurs between a switching neuron and its new network neurons, as a complement to the modulation of intrinsic properties that enable a switching neuron to exhibit dual frequency bursting.

In the STNS of the crab, *Cancer borealis*, activation of the modulatory commissural neuron 5 (MCN5) causes the pyloric-only LPG neuron to switch into dual pyloric- and gastric mill-timed bursting (12, 21). The LPG switch into gastric mill bursting occurs due to the MCN5 neuropeptide Gly^1^-SIFamide modulating a hyperpolarization-activated current, a persistent sodium current, and calcium, or calcium-related current(s) (12, 22). As part of our investigation into how the MCN5-elicited rhythm is generated, we found that LPG gastric mill-timed (slow) bursting is regulated by gastric mill network neurons, and that LPG can contribute to coordination among gastric mill network neurons (21), suggesting bidirectional communication between the LPG switching neuron and at least some of the gastric mill network neurons. However, in unmodulated saline conditions synapses between LPG and LG, IC, and DG gastric mill neurons have not been described. Therefore, we used two-electrode voltage clamp recordings to investigate whether there are functional synapses in the control saline condition, and whether the neuropeptide Gly^1^-SIFamide modulates synaptic transmission between LPG and LG, IC, and DG neurons, in addition to its established modulation of LPG’s intrinsic properties. We further tested for connectivity among the neurons by assessing the ability of LPG, LG, IC, and DG to entrain each other during Gly^1^-SIFamide application.

## Materials and methods

### Animals

Male *Cancer borealis* crabs were obtained from The Fresh Lobster company (Gloucester, MA) and maintained in tanks containing artificial seawater at 10-12 °C. For experiments, crabs were anesthetized on ice for 30-50 min, then during gross dissection, the foregut was removed from the animal, bisected, and pinned flat in a Slygard-170-lined dish (Thermo Fisher Scientific). During fine dissection, the stomatogastric nervous system (STNS) was dissected from the foregut and pinned in a Sylgard 184-lined petri dish (Thermo Fisher Scientific) (23). During the dissection, the preparation was kept chilled in *C. borealis* physiological saline at 4 °C.

### Solutions

*C. borealis* physiological saline was composed of the following (in mM): 440 NaCl, 26 MgCl_2_,13 CaCl_2_,11 KCl,10 Trizma base, 5 Maleic acid, pH 7.4-7.6. Squid internal electrode solution contained the following (in mM): 10 MgCl_2_, 400 Potassium D-gluconic acid, 10 HEPES, 15 Na_2_SO_4_, 20 NaCl, pH 7.45 (24). Gly^1^-SIFamide (GYRKPPFNG-SIFamide, custom peptide synthesis: Genscript) (12, 25–28) was dissolved in optima water (Thermo Fisher Scientific) at 10^−2^ M and stored at −20 °C. Tetrodotoxin (TTX) powder was dissolved in optima water at a final concentration of 10^−4^ M and stored at −20 °C. Stock solution of picrotoxin (PTX) was made by adding PTX powder (Sigma Aldrich) to 95% ethanol at 10^−2^ M and storing at −20 °C. For experiments, each substance was thawed and diluted in physiological saline at a final concentration of: Gly^1^-SIFamide, 5 µM; TTX, 1 µM and 0.1 µM; PTX, 10 µM.

### Electrophysiology

All preparations were continuously superfused with chilled *C. borealis* saline (8-10 °C), or chilled saline containing Gly^1^-SIFamide, PTX, and/or TTX, as indicated. For uninterrupted superfusion of the preparation, all solution changes were performed using a switching manifold. Extracellular nerve activity was recorded using a model 1700 A-M Systems Amplifier and custom-made stainless-steel pin electrodes. Vaseline wells were built around nerves to isolate neuron signals, with one stainless-steel electrode wire placed inside the well, and the other outside the well as reference. Stomatogastric ganglion (STG) somata were exposed by removing the thin layer of tissue across the ganglion and observed with light transmitted through a dark-field condenser (MBL-12010 Nikon Instruments). Intracellular recordings were collected using sharp-tip glass microelectrodes (18 – 40 MΩ) filled with squid internal electrode solution for entrainment experiments or filled with 0.6 M K_2_SO_4_ plus 20 mM KCl electrode solution for voltage clamp experiments. STG neurons and their locations were identified based on their intracellular activity, nerve projection patterns, and their interactions with other STG neurons. All intracellular recordings were collected using AxoClamp 900A amplifiers in current-clamp mode for entrainment experiments or two-electrode voltage clamp (TEVC)-mode for synapse experiments (Molecular Devices). All experiments were conducted in the isolated STNS following transection of the inferior and superior oesophageal nerves (*ion* and *son*, respectively). Electrophysiology recordings were collected using data acquisition hardware (Entrainment experiments: Micro1401, Cambridge Electronic Design; Synapse experiments: DigiData 1440A, Molecular Devices), software (Entrainment experiments: Spike2, ∼5 kHz sampling rate, Cambridge Electronic Design; Synapse experiments: Clampex 10.7, Molecular Devices, ∼5 kHz sampling rate) and laboratory computer (Dell).

Prior to all conducted experiments, unless otherwise indicated, the LP neuron was photoinactivated for the established bath-application model of MCN5 actions (12). Briefly, the LP neuron was impaled with a sharp microelectrode (30-40 MΩ) that was tip-filled with AlexaFluor-568 hydrazide (10 mM in 200 mM KCl, Thermo Fisher Scientific) and back-filled with squid internal solution (see *Solutions*). The LP soma was filled using constant hyperpolarizing current injection (−5 nA) for 5-10 min, and the current injection removed to allow the dye to diffuse to the neurites and axon in the *dvn* for 20 – 40 min. The STG was then illuminated for 5 – 7 min using a Texas red filter set (560 ± 40 nm wavelength; Leica Microsystems). Complete photoinactivation was confirmed when the LP membrane potential reached 0 mV.

### Graded synapses via two-electrode voltage clamp

To determine whether graded synapses between LPG and gastric mill neurons were present and/or modulated by Gly^1^-SIFamide, we used two-electrode voltage clamp (TEVC) recordings. Based on previous studies, we only proceeded with experiments and analyzed data when pre- and post-synaptic cells had an input resistance of at least 10 MΩ (24, 29). Action potentials were blocked with TTX (initial concentration: 1 µM, ∼5-10 min; final maintained concentration: 0.1 µM). When IC, LG, or DG was the presynaptic neuron, they were voltage clamped at −60 mV. The presynaptic neuron was stepped from its holding potential to −55 to −20 mV in 5 mV increments (5 s duration, 11 s inter-step interval). The postsynaptic cell was voltage clamped at −40 mV. For IC, LG, and DG to LPG synapses, the voltage step protocol was repeated in Gly^1^-SIFamide (SIF, 5 µM) plus TTX (0.1 µM) (TTX:SIF), following ∼10 min wash-in of TTX:SIF. To measure potential LPG to IC, LG, or DG synapses, the same voltage step protocol was used, however, PTX (10 µM) was used to block glutamatergic inhibitory synapses. LPG uses acetylcholine as its transmitter, and thus its actions are not blocked by PTX (30–32), enabling us to examine synapses from LPG onto LG, IC, and DG more directly. When LPG was the presynaptic neuron, it was held at −80 mV to turn off spontaneous activity of the other non-clamped LPG neuron. In TTX:PTX:Saline LPG was stepped from its holding potential to −55 mV to −20 mV in 5 mV increments, and the postsynaptic neuron current recorded with cells at a holding potential of −40 mV. This protocol was repeated in Gly^1^-SIFamide in TTX plus PTX (TTX:PTX:SIF), following ∼10 min wash-in of TTX:PTX:SIF. In a small subset of experiments as indicated, each presynaptic neuron (LG, IC, DG, or LPG), was clamped at a holding potential of −80 mV and stepped from −70 mV to −20 mV in 5 mV increments. Between each experimental condition, all neurons were held at −60 mV, near resting potential for STG neurons (33). Due to a factor of 10 error in a gain setting in TEVC in one amplifier, current amplitudes were multiplied post-hoc by a factor of 10. Before any adjustments were made, input resistance measurements in current clamp mode, unaffected by the gain error, from pre- and post-TEVC mode were compared to input resistance measurements in TEVC mode to verify the gain factor error. Additionally, the same neuron (e.g., LG) was not always recorded using the channel with the gain error in different preparations. Adjusted data matched the data recorded with the correct conversion factor in other experiments.

Voltage step protocols were repeated five times in each direction during each pair of synapses tested. Recordings were averaged and filtered using Clampfit 10.7 (pClamp, Molecular Devices, San Jose, CA) prior to analysis. Synaptic currents were measured by subtracting the mean holding current during the 2 s preceding each voltage step (Baseline) from the peak current within the first 2 s of the voltage step (Early) and the last 2 s of the voltage step (Late). For LPG to IC and to LG, we observed an inward current within the first 2 s of each voltage step. Thus, we also measured the peak amplitude of this inward current and subtracted it from the mean holding current.

### Gastric mill network entrainment

To examine whether each gastric mill-timed network neuron entrained each other, entrainment neurons (LPG [two copies], LG, IC, and DG [one copy each]) were stimulated with rhythmic depolarizing current injections (5 s pulse duration, 0.5 to 6 nA) at different entrainment frequencies (0.03 – 0.12 Hz; 5 s pulse, 6 to 28 s inter-pulse interval). Entrainment frequency started at 0.3 Hz or the first frequency (in 0.1 Hz increments) at which a neuron was able to be entrained. The amount of current injected into the entrainment neuron was chosen to match the firing frequency (± 5 Hz) of that neuron at steady state Gly^1^-SIFamide application prior to initiating entraining depolarizations. In some preparations, the entraining neuron was not active in Gly^1^-SIFamide, and we matched its firing frequency to the entraining neuron mean firing frequency from a previous data set (21). All entrainment manipulations were conducted during the last 20 min of a 30 min Gly^1^-SIFamide (5 µM) application. To examine LPG entrainment of gastric mill neurons LG, IC, and DG more directly, we performed an additional set of experiments in which picrotoxin (PTX, 10 µM), which blocks LG, IC, and DG glutamatergic inhibitory synapses, but not cholinergic LPG synapses (30–32), was added to the saline and Gly^1^-SIFamide (SIF:PTX) solutions. This condition isolates LPG from gastric mill input and blocks chemical synapses among the LG, IC, DG, neurons (30–32).

For each entrainment frequency, either the single LG, IC, or DG neuron, or both LPG neurons were rhythmically depolarized for ten cycles (11 bursts), and the instantaneous gastric mill burst frequency (1/cycle period [time between the first action potentials of two consecutive bursts]) for each test neuron (LG, IC, DG, and LPG) was measured. For LPG and IC, which can participate in both the pyloric and gastric mill rhythms, only gastric mill-timed bursts (defined below in Gastric mill burst identification) were used in analyses.

### Integer coupling analysis

To quantify the extent of entrainment between LG and IC, DG, and LPG, we used integer coupling as a measure of whether there was a fixed relationship between the instantaneous burst frequencies of each pair of neurons (34, 35). Integer coupling was measured by dividing the entraining neuron stimulation frequency by the instantaneous test neuron burst frequency and extracting the significand (the value to the right of the decimal point) from each ratio at each entrainment frequency, across preparations. The significand indicates a measure of how close a ratio is to an integer. For instance, if the ratio between the entrainment and instantaneous burst frequencies is 1.01, then significand is 0.01 and close to zero. If the ratio is 1.99, then the significand is 0.99 and close to 1. The cumulative probability distribution frequency (c.d.f.) of all significands for each entrainment frequency was calculated to determine how close each set of ratios was to an integer and plotted on a histogram. The shaded regions indicate the c.d.f. curves between entraining and test neurons at 0.06 Hz entrainment frequency (Figs. 6D, 7-8Ai, Bi). The blue dotted line indicates one extreme of perfect integer coupling, where all significands from all ratios are near 0 or 1, and the c.d.f. accumulates primarily at 0 or 1 (Figs. 6D, 7-8Ai, Bi) (35). On the other hand, if all significands across a group of data points are randomly and uniformly distributed between 0 and 1, then the c.d.f. of these values would be close to a diagonal line, and thus indicate zero integer coupling between neurons (solid red y = x line, Figs. 6D, 7-8Ai, Bi) (35).

To quantify the extent of integer coupling at each entrainment frequency, that is, the distance between the cumulative distribution and a random distribution (i.e., a diagonal line), we calculated the area between each c.d.f. curve and the diagonal line using a custom-written MATLAB script (MathWorks). For instance, if there was perfect integer coupling at a particular entrainment frequency, then the area between a c.d.f. falling along the blue dotted line and the diagonal would be 0.25, while an area close to zero would suggest there is no integer coupling between the entrainment and test neurons. To determine whether significands from a group of entrainment frequencies were significantly different from the diagonal line, we formulated a uniform distribution that would equal the y = x diagonal line, and thus have an area equal to zero to indicate no integer coupling. Then, for each entrainment frequency, we compared the experimental data to the uniform distribution using a two-sample Kolmogorov-Smirnov test (35). We used a two-sample test instead of a one-sample test because the calculated significands for the experimental data cannot be expected to be normally distributed (35). We divided the significance level by the number of entrainment frequencies (7) to correct for multiple comparisons. Thus, our significance threshold was 0.05/7 = 0.007. Entrainment frequencies with data from only one preparation were excluded from the integer coupling analysis.

The integer coupling analysis assumes test neuron integer-multiple relationships to the entrainment neuron (e.g., 1:1, 2:1, 3:1) using test neuron/entrainment neuron ratios. However, we did observe some fractional relationships in which there are fewer test neuron cycles relative to entrainment neuron (e.g., 1:2, 1:3; Fig. 6Biii). These entrainment relationships would result in significands between 0 and 1, for example a 1:2 entrainment would result in a significand of 0.5. The example “perfect” entrainment area of 0.25 (area between blue and red lines in figure 6D), represents equal proportions of significands occurring close to 0 or 1. Although the area calculation for “perfect” entrainment at intermediate significand values would differ from 0.25, it should still be greater than 0 and thus our area measurement enables us to quantify whether entrainment occurred, regardless of the type of entrainment.

### Gastric mill neuron burst identification

Only neurons that were bursting in gastric mill time during entrainment stimulations were included in the entrainment plots and integer coupling analysis. Criteria to determine gastric mill-timed bursting included: IC bursts had a duration of at least 0.45 s, with consecutive pyloric-timed IC bursts with burst durations of at least 0.45 s grouped together into a gastric mill burst, including up to one IC burst < 0.45 s in duration if it occurred between other IC bursts that were at least 0.45 s (21, 28); LG and DG bursts had at least 3 spikes per burst with a maximum 2 s interspike interval; and LPG gastric mill-timed bursts were identified as described (12). Briefly, a histogram containing LPG interspike intervals (ISIs) across an entire Gly^1^-SIFamide application was generated using a custom-written MATLAB script and the two largest peaks identified. The mean ISI between the first peak (∼0 – 0.5 s, intraburst intervals) and second peak (∼0.5 – 2 s, interburst intervals) was calculated and used as a cutoff value to identify LPG bursts, such that an ISI above the cutoff indicated the end of one burst and the beginning of another. A custom written Spike2 script that identifies LPG bursts with a duration greater than one pyloric cycle period (from PD burst onset to the next PD burst onset, ∼1 s) was then used to select only LPG gastric mill bursts from all LPG bursts identified with the ISI cutoff (12).

### Software and Statistics

Raw electrophysiology data were analyzed using Clampfit 10.7, as well as functions and scripts that were written in Spike2 and MATLAB. Graphs were plotted in SigmaPlot (Systat) or MATLAB and imported into CorelDraw (Corel). All final figures were created using CorelDraw. Statistical analyses were performed using SigmaPlot or MATLAB. Data were analyzed for normality using Shapiro-Wilk test before determining whether to use a parametric or nonparametric test on each dataset. A two-sample Kolmogorov-Smirnov test was used for integer coupling analysis. Two-way repeated measures (RM) analysis of variance (ANOVA) and Holm-Sidak post hoc tests for multiple comparisons were used for synaptic current analyses. Threshold for significance was p < 0.05, except for integer coupling analysis which had a threshold of p < 0.007. All data are presented as mean ± SEM.

### Code Accessibility

Scripts and functions used for analysis were written in Spike2 and/or MATLAB and are available at https://github.com/blitzdm/Fahoum_Blitz_2023_Synapses or upon request from D.M.B.

## Results

The crustacean stomatogastric nervous system (STNS) is a region of the central nervous system that controls feeding-related movements of the foregut, including the pyloric (food filtering, ∼1 Hz) and gastric mill (food chewing, ∼0.1 Hz) rhythms (36, 37) (Fig. 1A). The STNS is comprised of the paired commissural ganglia (CoG), single oesophageal ganglion (OG), the single stomatogastric ganglion (STG), and their connecting and peripheral nerves (Fig. 1A). The STG contains ∼30 neurons contributing to well-characterized CPG networks which drive the pyloric and gastric mill rhythms (Fig. 1B). Modulatory projection neurons originating in the CoGs and OG project to the STG where they alter intrinsic properties of individual neurons and/or the synapses between them to elicit multiple different versions of each rhythm (38–40).

**Figure 1.**
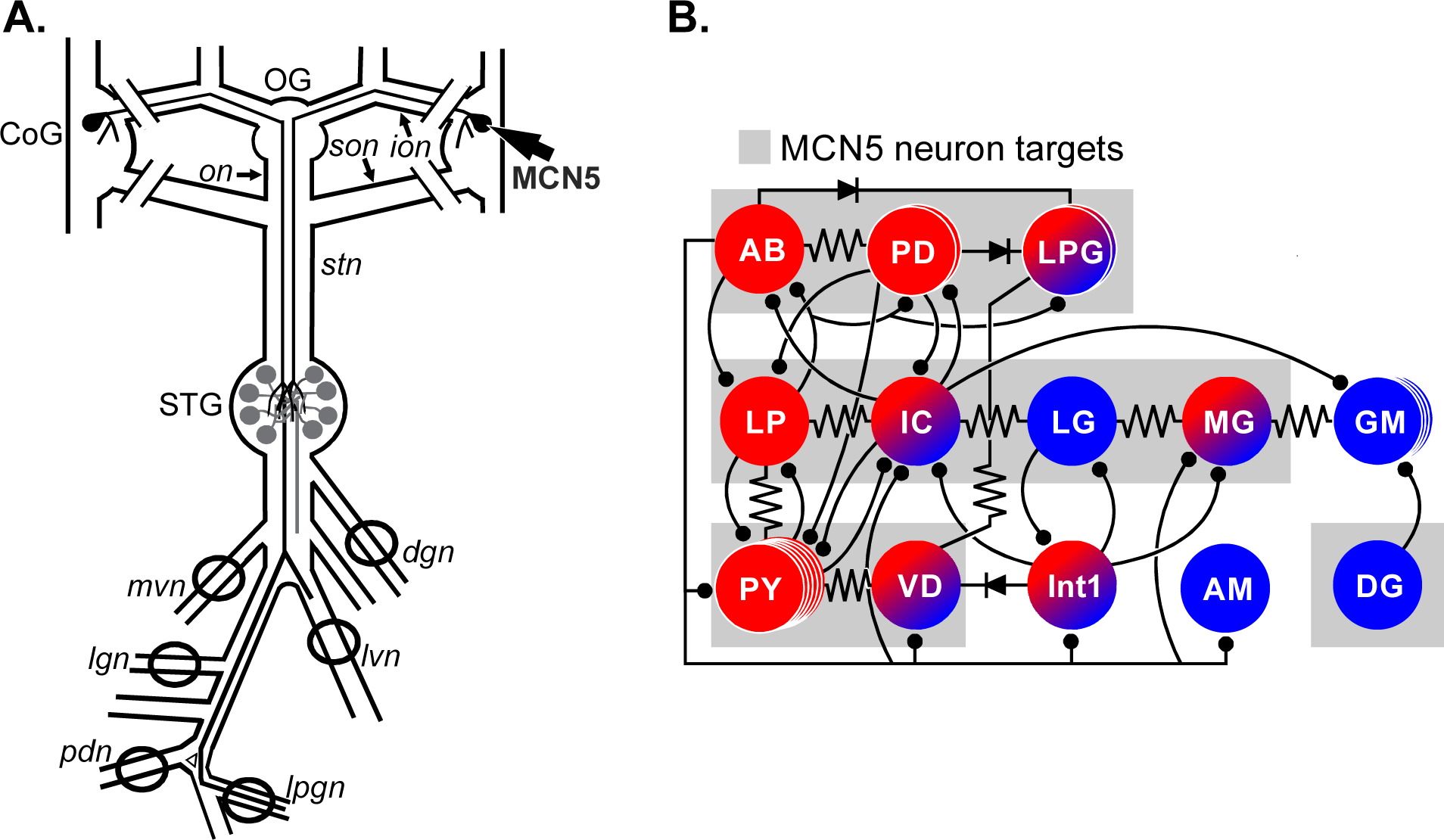
STNS schematic and circuit diagram of pyloric and gastric mill networks. ***A***, The modulatory projection neuron 5 (MCN5; two copies) has a soma (arrow) in each commissural ganglion (CoG) of the stomatogastric nervous system (STNS). The axon of each MCN5 neuron projects through the inferior oesophageal (*ion*), oesophageal (*on*), and stomatogastric (*stn*) nerves and terminates in the stomatogastric ganglion (STG) to synapse with motor neurons and interneurons responsible for generating rhythmic activity of the pyloric (∼1 Hz, food filtering) and gastric mill (∼0.1 Hz, food chewing) feeding-related behaviors. Each pyloric and gastric mill motor neuron has axons that project through posterior motor nerves to innervate muscles that control the movements for pyloric and gastric mill activity. Vaseline wells (*circles*) were built around motor nerves to record extracellular activity of these neurons. ***B***, The circuit diagram of the STG indicates known electrical (*resistor and diode symbols*) and inhibitory chemical (*ball and stick*) synapses among pyloric and gastric mill neurons in saline conditions. When active, the MCN5 neuron targets both pyloric and gastric mill neurons (*grey boxes*), however, synaptic modulation targets of MCN5 are unknown. Abbreviations, Ganglia: CoG, commissural ganglion; OG, oesophageal ganglion; STG, stomatogastric ganglion; Nerves: *dgn*, dorsal gastric nerve; *ion*, inferior oesophageal nerve; *lgn*, lateral gastric nerve; *lpgn* lateral posterior gastric nerve; *lvn*, lateral ventricular nerve; *mvn*, medial ventricular nerve; *on*, oesophageal nerve; *pdn*, pyloric dilator nerve; *son*, superior oesophageal nerve*; stn*, stomatogastric nerve; Neurons: AB, anterior burster; AM, anterior medial; DG, dorsal gastric; GM, gastric mill; IC, inferior cardiac; Int 1, interneuron 1; LG, lateral gastric; LP, lateral pyloric; LPG, lateral posterior gastric; MG, medial gastric; PD, pyloric dilator; PY, pyloric; VD, ventricular dilator.

The projection neuron modulatory commissural neuron 5 (MCN5) is located in the CoG and projects its axons to the STG via the inferior oesophageal and stomatogastric nerves (*ion* and *stn*, respectively; Fig. 1A). Bath application or neuronal release of the MCN5 neuropeptide Gly^1^-SIFamide increases pyloric frequency (41) and activates the gastric mill rhythm (12, 28). Furthermore, MCN5 activation or bath application of its neuropeptide Gly^1^-SIFamide (5 µM) switches the LPG neuron from pyloric-only activity to dual pyloric/gastric mill-timed bursting (fast/slow, respectively) (12, 21, 28). LPG is active in pyloric time due to electrical coupling to the other pyloric pacemaker neurons (AB, PD; Fig. 1B) and generates slower, gastric mill-timed bursts due to Gly^1^-SIFamide modulation of LPG intrinsic properties (12, 22). Although the LPG gastric mill-timed bursting does not require synaptic input from the gastric mill network, these oscillations are regulated by gastric mill network neurons including being coordinated with the rest of the gastric mill network. Further, LPG slow bursting regulates LG and dorsal gastric (DG) neuron activity, and LPG can coordinate gastric mill network neurons (21), suggesting the presence of synapses between LPG and the gastric mill neurons. Synapses between LPG and gastric mill network neurons have not been described (Fig. 1B) (42). We hypothesized that Gly^1^-SIFamide modulates synapses between LPG and LG, IC, and DG to become functional in this particular modulatory state.

### Gly^1^-SIFamide modulates synapses between LG and LPG

Among STG neurons, neurotransmitter release is primarily controlled in a graded, voltage-dependent manner by subthreshold membrane potential oscillations (43–46). Therefore, we used two-electrode voltage clamp (TEVC) recordings in pairs of neurons (LPG:LG, LPG:IC, and LPG:DG), in TTX (0.1 µM) to block action potentials, and examined whether a synaptic current was elicited in response to presynaptic voltage steps.

We first examined whether there was a graded synapse from LG to LPG. LG was voltage clamped at −60 mV and stepped from −55 mV to −20 mV in 5 mV increments (5 s duration, 11 s inter-step interval) and LPG was voltage clamped at −40 mV (Fig. 2A). In TTX:Saline, there was no apparent synaptic current elicited in LPG, suggesting that there was no functional synapse from LG to LPG in saline conditions (Fig. 2A, black traces). When TTX plus Gly^1^-SIFamide (TTX:SIF) was applied to the preparation, an outward current was evoked during LG voltage steps, with increased amplitude at more depolarized voltage steps (Fig. 2A, red traces). Additionally, we observed “bumps” in the LPG synaptic currents, such as in the grey trace, that were similar to transient currents that we observed between LG voltage steps (Fig. 2A, grey arrows, n = 4/4 preparations). We hypothesized that these “bumps” reflect synaptic input due to Gly^1^-SIFamide-elicited activity in the DG and/or IC neuron. However, the early outward current (during the “Early” bracket), which increased in amplitude with greater depolarization of the LG neuron, was consistently time-locked to the LG voltage steps, and is thus a synaptic current elicited by depolarization of the LG neuron (Fig. 2A, red traces). To quantify the LPG response to LG voltage steps, we subtracted the mean holding current (Baseline) across two seconds immediately prior to the voltage step from the peak amplitude of LPG current within the first two seconds of the voltage step (Early) and the last two seconds of the voltage step (Late) (Fig. 2A, blue brackets). LPG synaptic current was plotted as a function of LG voltage steps for each of the two measurement time points (Fig. 2Bi-Bii; TTX:Saline black symbols, TTX:SIF red symbols; each symbol represents one experiment). We used a two-way repeated measures (RM) ANOVA for statistical analysis with condition (TTX:Saline vs. TTX:SIF), and voltage as the two main effects. For the LPG synaptic current during the first 2 s, there was a significant interaction of condition and voltage (Fig. 2Bi) (F[3,7] = 2.92, p = 0.027, Two Way RM ANOVA; n = 4). Specifically, the peak current amplitude was different between TTX:Saline and TTX:SIF between −35 and −20 mV, except for −30 mV (Table 1; n = 4). As a graded synaptic current should express voltage-dependence, we used a comparison of the measured current between −55 and −20 mV for an indicator of voltage-dependence. There was no difference in current between these two voltages in the TTX:Saline condition, but there was a larger current at −20 mV compared to −55 mV in the TTX:SIF condition (Table 1; n = 4). During the last 2 s of each LG voltage step, there was also a significant interaction between condition and voltage on the LPG synaptic current (Fig. 2Bii) (F[3,7] = 5.01, p = 0.002, Two Way RM ANOVA, n = 4). There was a larger LPG synaptic current during Gly^1^-SIFamide compared to TTX:Saline during LG depolarizing steps from −35 to −20 mV (Table 1; n = 4). Similar to the first 2 s, there was no voltage-dependence in TTX:Saline during the last 2 s of the voltage steps, but there were differences in the synaptic current amplitudes between −55 and −20 mV during TTX:SIF (Table 1; n = 4). Thus, Gly^1^-SIFamide enhanced a synapse from LG to LPG, which was dependent upon presynaptic voltage, that was not evident in the TTX:Saline condition.

**Figure 2.**
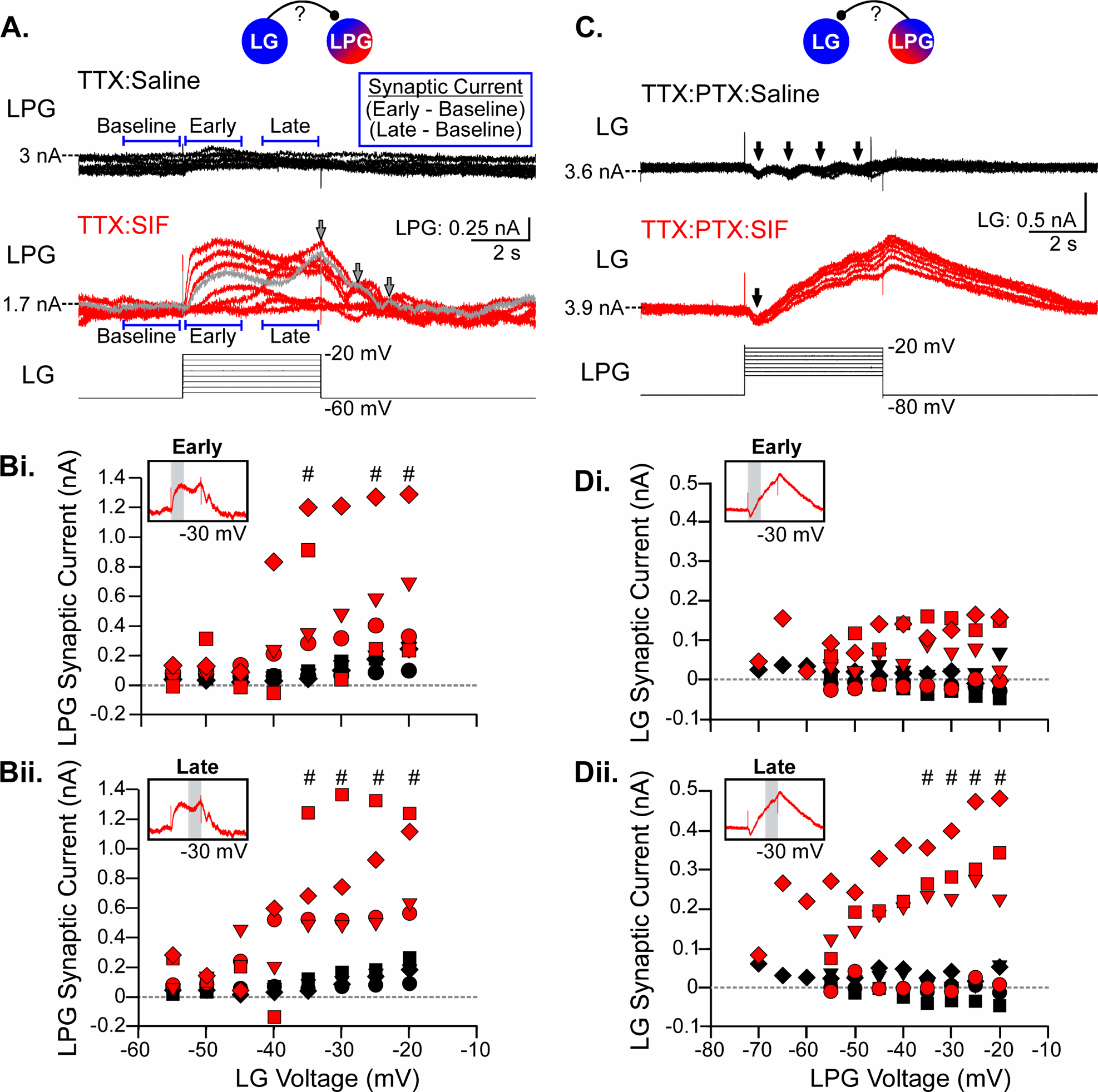
Gly^1^-SIFamide modulates graded synaptic transmission between LG and LPG. Intracellular two-electrode voltage clamp (TEVC) recordings were performed to examine synapses between LG and LPG. Symbols at the top indicate the synapse being tested in the left and right columns. ***A***, LPG was voltage clamped at −40 mV while LG was held at −60 mV and stepped from −55 mV to −20 mV in 5 mV increments (5 s duration, 11 s inter-step interval) in either TTX:Saline (***top***, black traces) or TTX:SIF (***bottom***, red traces). Grey arrows and grey trace indicate “bumps” in LPG voltage that occurred both during and between the LG voltage steps. Blue brackets indicate the time periods used for analysis. For each time point, the baseline current was subtracted from the peak current within that time window. ***Bi-Bii***, LPG synaptic current in TTX:Saline (black) and TTX:SIF (red) as a function of LG voltage steps during the Early (***Bi***) and Late (***Bii***) component of the voltage step. Different shapes in Bi-Bii and Di-Dii represent individual experiments (n = 4). Insets indicate timepoints of measurements during an example voltage step. ***C***, LPG was voltage clamped at −80 mV to eliminate spontaneous bursting activity of the other non-clamped LPG neuron between voltage steps and stepped to −55 mV to −20 mV in 5 mV increments (5 s duration, 11 s inter-step interval) with LG held at −40 mV. 10 µM PTX was used to block glutamatergic transmission. Synaptic currents were measured in TTX:PTX:Saline (***top*,** black traces) and TTX:PTX:SIF (***bottom***, red traces). ***Di-Dii***, LG synaptic current as a function of LPG voltage steps during the Early (***Di***) and Late components of the LPG voltage step. ^#^p < 0.05 TTX:Saline vs. TTX:SIF (Bi-ii) or TTX:PTX:Saline vs. TTX:PTX:SIF (Di-ii) at marked voltages (Two-way ANOVA, Holm-Sidak post hoc; see Table 1).

**Table 1.**
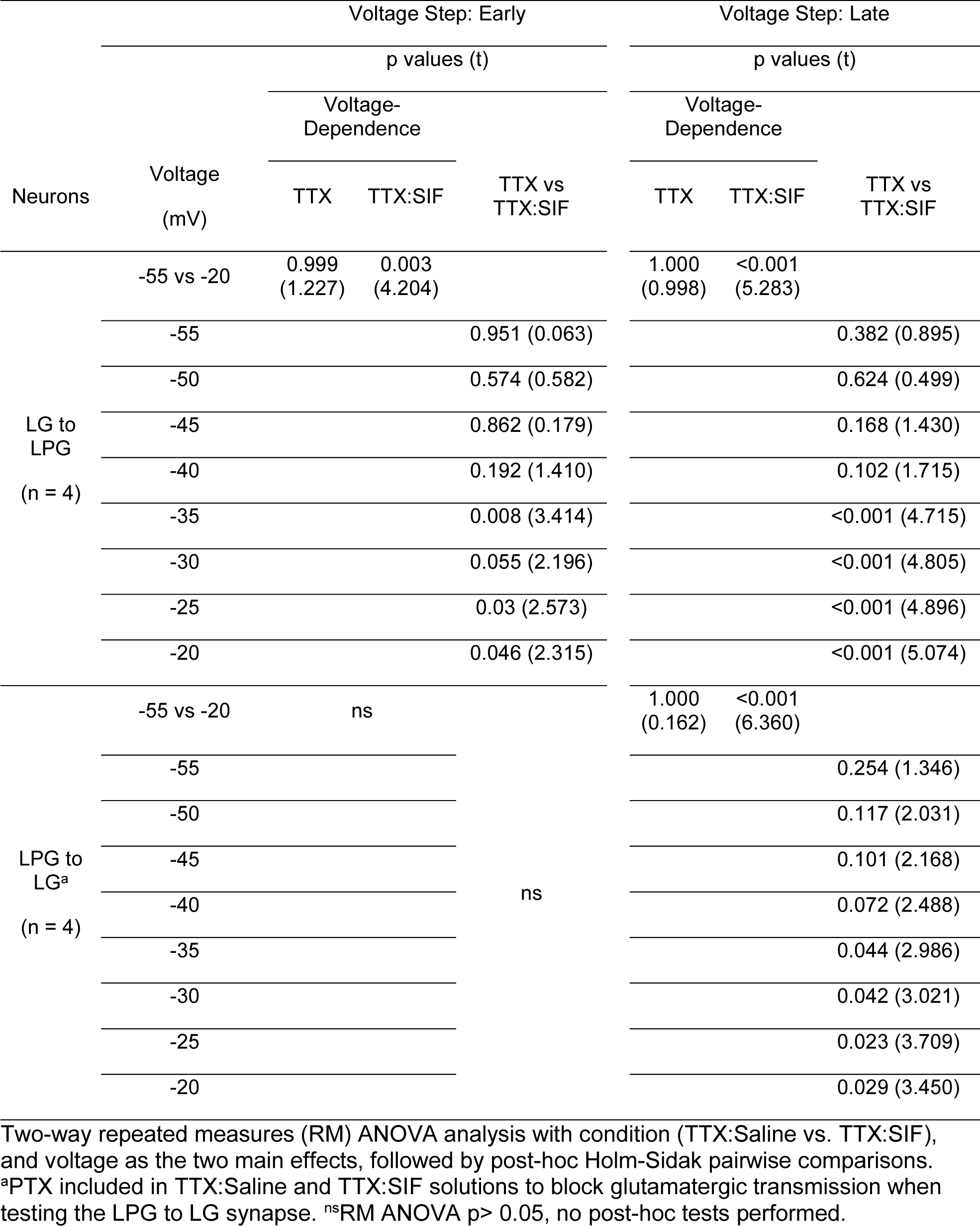
Effects of voltage and Gly^1^-SIFamide on synaptic currents between LG and LPG, at early and late time windows in the voltage step.

We next tested for a synaptic connection from LPG to LG. TTX was applied with picrotoxin (PTX, 1 µM) to block LG, IC, and DG inhibitory glutamatergic synapses to examine LPG cholinergic synapses onto LG without complications of chemical transmission from the other gastric mill neurons that were not voltage clamped. Since we performed TEVC with only one of the two LPG neurons, LPG was voltage clamped at −80 mV to eliminate spontaneous bursting activity of the other non-clamped LPG neuron between voltage steps. The voltage-clamped LPG neuron was stepped from its holding voltage to −55 to −20 mV in 5 mV increments, or in a subset of experiments to additional voltage steps ranging from −70 mV to −20 mV (5 s duration, 11 s inter-step interval) (Fig. 2C). LG was voltage clamped at −40 mV. In saline conditions, during LPG voltage steps, a series of small amplitude, inward currents were evoked in LG (Fig. 2C, black traces, black arrows). During application of Gly^1^-SIFamide, there was still an initial inward current in LG within the first two seconds of each LPG voltage step, but the small repetitive currents were then summed with a larger amplitude outward current (Fig. 2C, red traces). The outward current in LG increased with increasing LPG voltage steps, indicating graded transmitter release (Fig. 2C, red traces).

We quantified LG synaptic current from LPG in the same way as described above at the Early and Late time points, with LG synaptic current plotted as a function of LPG voltage steps (Fig. 2Di-Dii). For the peak LG synaptic current during the Early part of the voltage step, there was no significant interaction of condition and voltage (Fig. 2Di) (F[3,7] = 2.267, p = 0.069), nor a main effect of voltage (F[3,7] = 1.61, p = 0.187) or condition (F[3,1] = 3.95, p = 0.141; Two-way RM ANOVA; n = 4). However, during the Late part of each LPG voltage step, there was a significant interaction between condition and voltage (Fig. 2Dii) (F[3,7] = 5.11, p = 0.002, Two-way RM ANOVA; n = 4). Specifically, the peak LG synaptic current amplitude was different between TTX:Saline and TTX:SIF between −35 and −20 mV (Fig. 2Dii; Table 1; n = 4). Further, the synaptic current amplitudes were different between −55 and −20 mV in TTX:SIF (Table 1; n = 4) indicating voltage-dependence, but not during TTX:Saline (Table 1; n = 4). Thus, the LPG to LG current appeared to have a slower onset than the LG to LPG synaptic current as there was only a significant increase in current in the Late component of the LPG voltage step (Fig. 2Dii). In the STNS, cholinergic inhibition has a slower time course than glutamatergic inhibition (32, 47, 48), aligning with the apparent differences in timing between the LG- (glutamatergic) and LPG- (cholinergic) elicited synaptic currents.

Unlike the outward current elicited in LG, the inward current was not voltage dependent or modulated by Gly^1^-SIFamide. As the inward current summed with the outward current beyond the beginning of the voltage step, we only quantified its amplitude during the initial 2 s period. We found no interaction between condition and voltage (F[1,7] = 0.34, p = 0.911), and no effect of condition (F[1,1] = 3.25, p = 0.071) or voltage (F[1,7] = 0.71, p = 0.554; Two-way RM ANOVA, n = 4).

### Gly^1^-SIFamide modulates synapses between IC and LPG

We next examined whether a graded synapse was present from IC to LPG. Using the same protocol as above, in TTX:Saline, IC voltage steps elicited a small outward current during steps from −35 to −20 mV indicating a functional synapse between IC and LPG in Saline (Fig. 3A, black traces, n = 5/5). When TTX:SIF was applied to the preparation, a larger outward current was evoked during IC voltage steps, with increased amplitude at more depolarized voltage steps (Fig. 3A, red traces). Similar to the LPG synaptic current from LG voltage steps, we observed that the LPG outward synaptic current had a steep slope as it reached its peak in the first two seconds of IC voltage steps at the more depolarized voltages (Fig. 3A, red traces). In 5/5 preparations, we observed “bumps” in LPG synaptic current during IC voltage steps that appeared similar to those observed during LG voltage steps (Fig. 3A, red trace, grey arrows). However, these currents occurred only during IC voltage steps, not when IC was held at −60 mV.

**Figure 3.**
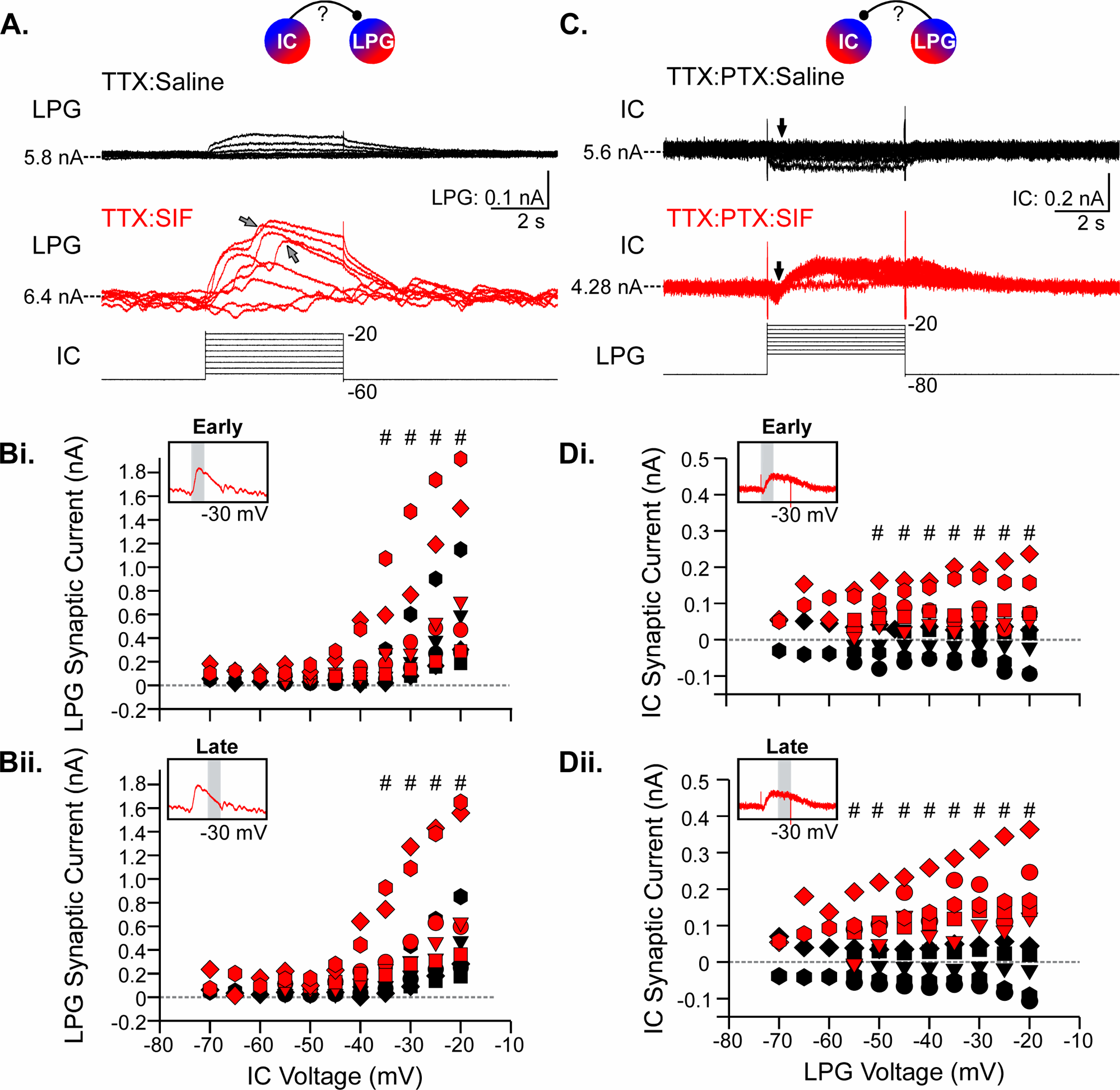
Gly^1^-SIFamide modulates graded synaptic transmission between IC and LPG. Intracellular TEVC recordings were used to examine graded synapses between IC and LPG. Symbols at the top indicate the synapse being tested in the left and right columns. ***A***, LPG voltage clamped at −40 mV and IC stepped from −60 mV to −55 mV to −20 mV in 5 mV increments in TTX:Saline (***top***, black traces) or TTX:SIF (***bottom***, red traces). Grey arrows indicate “bumps” in LPG voltage that occurred only during the IC voltage steps. ***Bi-Bii****, **Di-Dii**,* Same analysis as in figure 2A but for IC and LPG neurons (n = 5). ***C***, Example recordings of LPG held at −80 mV to eliminate spontaneous bursting activity of the other non-clamped LPG neuron between voltage steps and stepped to −55 mV to −20 mV with IC held at −40 mV. 10 µM PTX was used to block glutamatergic transmission. Synaptic currents were measured in TTX:PTX:Saline (***top*,** black traces) and TTX:PTX:SIF (***bottom***, red traces). ^#^p < 0.05 TTX:Saline vs. TTX:SIF (Bi-ii) or TTX:PTX:Saline vs. TTX:PTX:SIF (Di-ii) at marked voltages (Two-way ANOVA, Holm-Sidak post hoc; see Table 2).

**Table 2.**
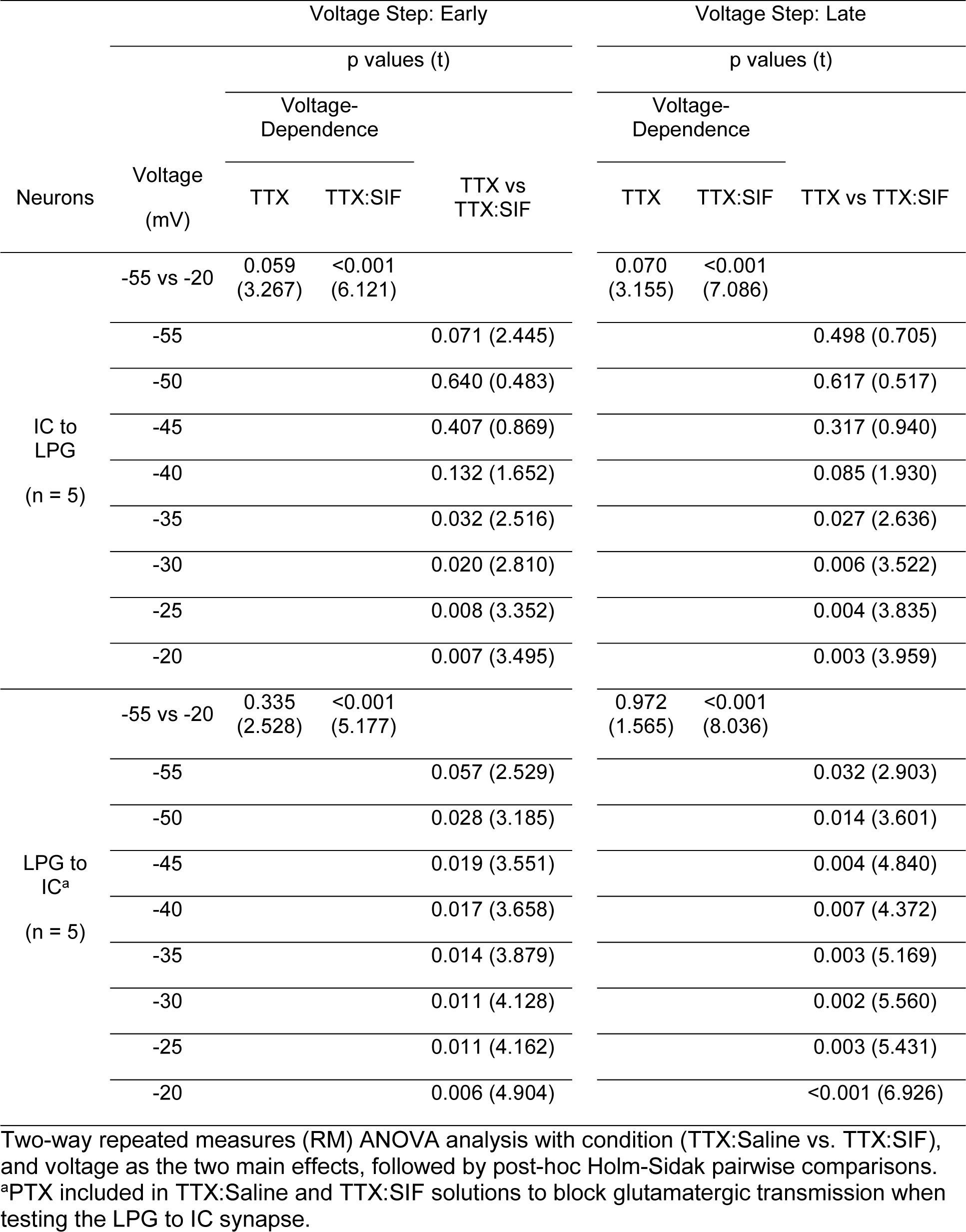
Effects of voltage and Gly^1^-SIFamide on synaptic currents between IC and LPG, at early and late time windows in the voltage step.

Across preparations, during the Early time point of the LPG synaptic current, there was a significant interaction of condition and voltage (Fig. 3Bi) (F[4,7] = 3.81, p = 0.005; Two-way RM ANOVA; n = 5). Specifically, the peak LPG current amplitude was different between TTX:Saline and TTX:SIF between −35 and −20 mV (Table 2; n = 5). In addition, there was a difference in the LPG synaptic current amplitude between −55 mV and −20 mV in TTX:SIF, but not during TTX:Saline (Table 2; n = 5). During the Late time window of each IC voltage step, there was also a significant interaction between condition and voltage on the LPG synaptic current (Fig. 3Bii) (F(4, 7) = 5.07, p < 0.001; Two-way RM ANOVA; n = 5). LPG synaptic current was larger during TTX:SIF compared to TTX:Saline during IC depolarizing voltage steps between −35 and −20 mV (Table 2; n = 5). Similar to the Early component, there was no voltage dependence in TTX:Saline during the Late component of the IC voltage steps, but the LPG synaptic current amplitude was different between −55 mV and −20 mV during TTX:SIF (Table 2; n = 5). Thus, there were synaptic currents present in the Saline condition, and Gly^1^-SIFamide enhanced this synapse from IC to LPG.

We also found a modulated synapse in the opposite direction, from LPG to IC. As explained above, LPG was voltage clamped at −80 mV and stepped from −70 to −20 mV or from −55 to −20 mV in 5 mV increments (Fig. 3C-D), with PTX included to block glutamatergic transmission. IC was voltage clamped at −40 mV. In saline conditions, IC responded to LPG voltage steps by generating a small amplitude, downward peak (Fig. 3C, black traces, black arrows, n = 4/5). During Gly^1^-SIFamide plus TTX:PTX application, there was still an initial downward response (inward, excitatory current, n = 5/5) in the first two seconds of each LPG voltage step with a subsequent outward inhibitory current in IC that increased with more depolarized LPG voltage steps, suggesting that LPG inhibited IC in a graded manner (Fig. 3C, red traces).

Across experiments, there was an increased amplitude of LPG-elicited synaptic current in IC, with a presynaptic voltage-dependence, in Gly^1^-SIFamide. For the IC synaptic current in the Early component of each LPG voltage step, there was a significant interaction of condition and voltage (Fig. 3Di) (F[4,7] = 6.57, p < 0.001; Two-way RM ANOVA; n = 5). The Early IC synaptic current amplitude was different between TTX:PTX:Saline and TTX:PTX:SIF from −50 to −20 mV (Table 2; n = 5). Furthermore, there was a difference in the IC current amplitude at −55 vs. −20 mV in TTX:PTX:SIF, but not during TTX:PTX:Saline (Table 2; n = 5). During the Late component of LPG voltage steps, there was also a significant interaction between condition and voltage (Fig. 3Dii) (F[4,7] = 10.75, p < 0.001; Two-way RM ANOVA; n = 5). There was a larger IC synaptic current during TTX:PTX:SIF compared to TTX:PTX:Saline during LPG depolarizing voltage steps from −55 to −20 mV (Table 2; n = 5). Furthermore, IC synaptic current amplitude during the Late component displayed voltage dependence during TTX:PTX:SIF application, but not during TTX:PTX:Saline (−55 vs −20 mV; Table 2; n = 5).

We also measured the peak inward current in IC in saline and Gly^1^-SIFamide conditions. We found an effect of condition (F[4,1] = 11.85, p = 0.026) and LPG voltage (F[4,7] = 7.13, p < 0.001), but no interaction between condition and voltage (F[4,7] = 1.28, p = 0.299; Two-way RM ANOVA; n = 5). In conclusion, these results suggest that a voltage dependent inhibitory graded synapse from LPG onto IC was not functional in baseline condition but was enhanced by Gly^1^-SIFamide. For this pair, there was little difference in the Early vs. Late components, however in the example traces (Fig. 3C), the rise time of the IC synaptic current does appear slower than the rise time of the IC to LPG synapse (Fig. 3A).

### Gly^1^-SIFamide modulates synapses between DG and LPG

For the last pair of neurons, DG and LPG, there was no apparent synaptic current elicited in LPG in an example experiment in TTX:Saline (Fig. 4A, black traces). When TTX:SIF was applied to the preparation, a small outward current was evoked during DG voltage steps, but the amplitude did not appear to change with more depolarized voltage steps (Fig. 4A, red traces). Although small, at more depolarized DG voltages (−25 to −20 mV), the LPG outward synaptic current reached its peak rapidly (Fig. 4A, red traces), compared to the example LPG to DG synaptic currents (Fig. 4C).

**Figure 4.**
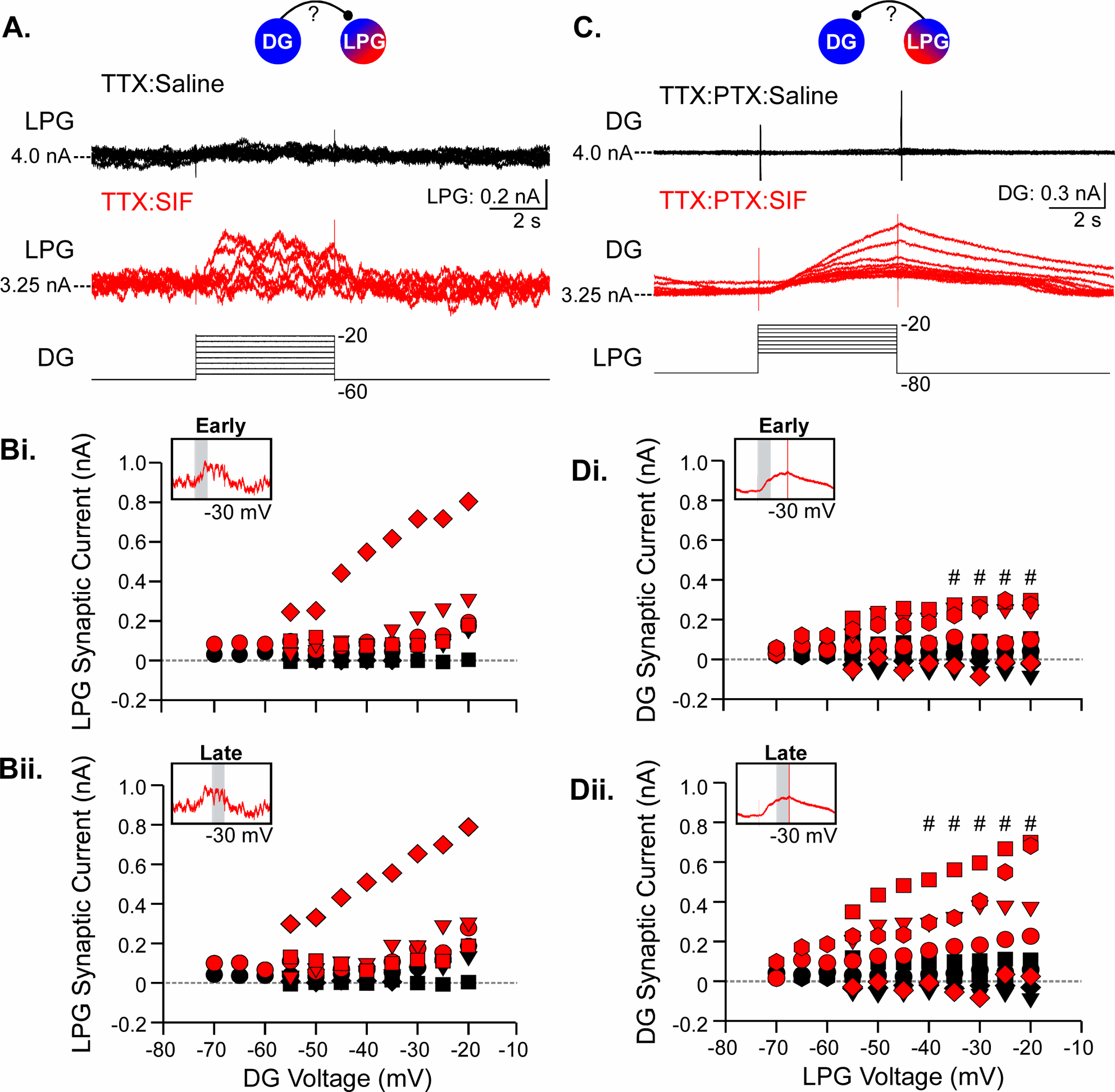
Gly^1^-SIFamide modulates graded synaptic transmission from LPG to DG, but not DG to LPG. Intracellular TEVC recordings were performed to examine synapses between DG and LPG. ***A***, Example recordings of LPG (voltage clamped at −40 mV) and DG held at −60 mV and stepped from −55 mV to −20 mV in 5 mV increments in TTX:Saline (***top***, black traces) or TTX:SIF (***bottom***, red traces). LPG synaptic current as a function of DG voltage steps is plotted for Early (***Bi***) and Late (***Bii***) timepoints (n=4). ***C***, Example recordings of LPG voltage clamped at −80 mV and stepped to −55 mV to −20 mV with DG held at −40 mV. Synaptic currents were measured in TTX:PTX:Saline (***top*,** black traces) and TTX:PTX:SIF (***bottom***, red traces, transient in DG current trace truncated for presentation purposes). ***Di-Dii,*** DG synaptic current as a function of LPG voltage steps during Early (***Di***) and Late time points (n=5). ^#^p < 0.05 TTX:Saline vs. TTX:SIF (Bi-ii) or TTX:PTX:Saline vs. TTX:PTX:SIF (Di-ii) at marked voltages (Two-way ANOVA, Holm-Sidak post hoc; see Table 3).

**Table 3.** Effects of voltage and Gly^1^-SIFamide on synaptic currents between DG and LPG, at early and late time windows in the voltage step.

| Neurons | Voltage Step: Early |  |  |  | Voltage Step: Late |  |  |
| --- | --- | --- | --- | --- | --- | --- | --- |
|  | Voltage<br>(mV) | p values (t) |  |  | p values (t) |  |  |
|  |  | Voltage-Dependence |  |  | Voltage-Dependence |  |  |
|  |  | TTX | TTX:SIF | TTX vs TTX:SIF | TTX | TTX:SIF | TTX vs TTX:SIF |
| DG to LPG<br>(n = 4) | -55 vs -20 | 0.415<br>(2.459) | <0.001<br>(4.996) |  | 0.190<br>(2.848) | <0.001<br>(6.262) |  |
|  | -55 |  |  | 0.294 (1.187) |  |  | 0.207 (1.521) |
|  | -50 |  |  | 0.274 (1.244) |  |  | 0.205 (1.528) |
|  | -45 |  |  | 0.139 (1.793) |  |  | 0.137 (1.883) |
|  | -40 |  |  | 0.101 (2.054) |  |  | 0.136 (1.885) |
|  | -35 |  |  | 0.064 (2.441) |  |  | 0.064 (2.593) |
|  | -30 |  |  | 0.134 (1.826) |  |  | 0.104 (2.126) |
|  | -25 |  |  | 0.137 (1.804) |  |  | 0.099 (2.178) |
|  | -20 |  |  | 0.103 (2.036) |  |  | 0.072 (2.476) |
| LPG to DG<sup>a</sup><br>(n = 5) | -55 vs -20 | 1.000<br>(0.156) | <0.001<br>(6.058) |  | 1.000<br>(0.128) | <0.001<br>(7.501) |  |
|  | -55 |  |  | 0.177 (1.605) |  |  | 0.131 (1.807) |
|  | -50 |  |  | 0.073 (2.342) |  |  | 0.065 (2.363) |
|  | -45 |  |  | 0.100 (2.072) |  |  | 0.062 (2.396) |
|  | -40 |  |  | 0.078 (2.283) |  |  | 0.043 (2.700) |
|  | -35 |  |  | 0.043 (2.816) |  |  | 0.034 (2.902) |
|  | -30 |  |  | 0.057 (2.556) |  |  | 0.024 (3.199) |
|  | -25 |  |  | 0.033 (3.066) |  |  | 0.010 (4.042) |
|  | -20 |  |  | 0.035 (3.003) |  |  | 0.007 (4.413) |

Two-way repeated measures (RM) ANOVA analysis with condition (TTX:Saline vs. TTX:SIF), and voltage as the two main effects, followed by post-hoc Holm-Sidak pairwise comparisons. <sup>a</sup>PTX included in TTX:Saline and TTX:SIF solutions to block glutamatergic transmission when testing the LPG to DG synapse.

With one exception, DG to LPG synaptic currents were small relative to those elicited by IC and LG voltages (Figs. 2-4). Using the same quantification of synaptic currents as above, we found for the Early LPG synaptic current, there was only an effect of voltage (Fig. 4Bi) (F[3, 7] = 4.86, p = 0.002, Two-way RM ANOVA; n = 4), such that the LPG synaptic current amplitude was different between −55 vs. −20 mV during TTX:SIF, but not during TTX:Saline (Table 3; n = 4). There was no effect of condition (F[3, 1] = 4.58, p = 0.122), nor interaction between condition and voltage on LPG current during the Early component (F[3, 7] = 0.98, p = 0.475; Two-way RM ANOVA; n = 4). Similarly, during the Late component of each DG voltage step, there was an effect of voltage (Fig. 4Bii) (F[3,7] = 6.79, p < 0.001; Two-way RM ANOVA; n = 4), such that there were differences in LPG synaptic current amplitudes between DG voltage steps to −55 vs. −20 mV in TTX:SIF, but not during TTX:Saline (Table 3; n = 4). However, there was no effect of condition (F[3, 1] = 5.34, p = 0.104), nor an interaction between condition and voltage (F[3, 7] = 1.88, p = 0.132; Two-way RM ANOVA; n = 4). Overall, although there was no statistical difference between saline and Gly^1^-SIFamide conditions, there was a voltage-dependent effect on LPG current only during Gly^1^-SIFamide. This suggests that the Gly^1^-SIFamide peptide did modulate a DG-to-LPG synapse that was not effective at baseline, however the peptide effects were quite variable. The effects on the DG to LPG synaptic currents included a quadrupling of current amplitude (n=1/4), no change in amplitude (n = 1/4) and a small but evident increase in strength (n = 2/4; Fig. 4A, Bi). We did not explore this variability further.

Lastly, we tested whether a graded synapse from LPG to DG was present. In TTX:PTX:Saline, DG current did not appear to change relative to baseline current, suggesting that there was no functional synapse between LPG and DG in saline conditions (Fig. 4C, black traces). During Gly^1^-SIFamide application, DG generated an outward inhibitory current that increased with increasing LPG voltage steps, suggesting that LPG inhibited DG in a graded manner (Fig. 4C, red traces). As in LG and IC, the LPG-elicited synaptic current in DG had a slower time-course than the DG to LPG synaptic current.

Quantification and statistical analysis across experiments supported modulation of an LPG to DG synapses as in the example experiment. For the Early component of the DG synaptic current, there was a significant interaction of condition and voltage (Fig. 4Di) (F[4, 7] = 4.45, p = 0.002; Two-way RM ANOVA; n = 5). Specifically, the Early DG synaptic current amplitude was different between TTX:PTX:Saline and TTX:PTX:SIF between −35 and −20 mV, except for −30 mV (Table 3; n = 5). Furthermore, there was a difference in the DG synaptic current amplitude at −55 vs. −20 mV in TTX:PTX:SIF, but not during TTX:PTX:Saline (Table 3; n = 5). During the Late component of each LPG voltage step, there was also a significant interaction of condition and voltage (Fig. 4Dii) (F[4, 7] = 6.49, p < 0.001; Two-way RM ANOVA; n = 5). There was a larger DG synaptic current during TTX:PTX:SIF compared to TTX:PTX:Saline during LPG depolarizing steps from −40 to −20 mV (Table 3; n = 5). Additionally, voltage dependence of the DG synaptic current amplitudes was evident during TTX:PTX:SIF, but not during TTX:PTX:Saline (−55 vs. −20 mV; Table 3; n = 5). Thus, Gly^1^-SIFamide enhanced a synapse from LPG to DG, which was dependent upon presynaptic voltage, that was not functional in the TTX:PTX:Saline condition.

Collectively, our voltage clamp experiments indicated that there are graded synapses, which are primarily ineffective in TTX:Saline, that are modulated to become effective, or to become stronger (IC to LPG synapse), during Gly^1^-SIFamide application. In some modulatory conditions, graded transmission can be sufficient for CPG neurons to generate coordinated network oscillations in the absence of spike-mediated transmission (43, 46, 49). We tested whether graded transmission is also sufficient to drive the Gly^1^-SIFamide gastric mill rhythm in the absence of action potentials.

In normal saline conditions, recording from the neurons in current-clamp mode, only the pyloric rhythm was active, with LPG and PD (*pdn* recording) generating pyloric-timed bursting activity (Fig. 5A, left). During application of Gly^1^-SIFamide (5 µM), there was a coordinated gastric mill rhythm between LPG, LG, IC, and DG neurons (Fig. 5A, right) (12, 21, 28). When spike-mediated transmission was blocked by eliminating action potential generation with TTX (0.1 µM) (Fig. 5B), the neurons exhibited stable resting membrane potentials without oscillations. However, during TTX plus Gly^1^-SIFamide application (5 µM), LPG, LG, IC, and DG generated coordinated gastric mill-timed oscillations (Fig. 5B, right; n = 2). There was coordination of both pyloric (grey lines) and gastric mill-timed (e.g., shaded box) oscillations between the neurons (Fig. 5B, right). Inhibitory relationships are suggested, for example, in the most hyperpolarized membrane potentials in DG occurring during the most depolarized membrane potentials in LPG (Fig. 5B, dotted box). Thus, graded transmission was sufficient to mediate coordinated gastric mill network activity in the Gly^1^-SIFamide modulatory state. These experiments plus our previous study (21) demonstrated that LPG and the LG, IC, DG group of gastric mill neurons contribute to coordination of gastric mill bursting in the Gly^1^-SIFamide modulatory state.

**Figure 5.**
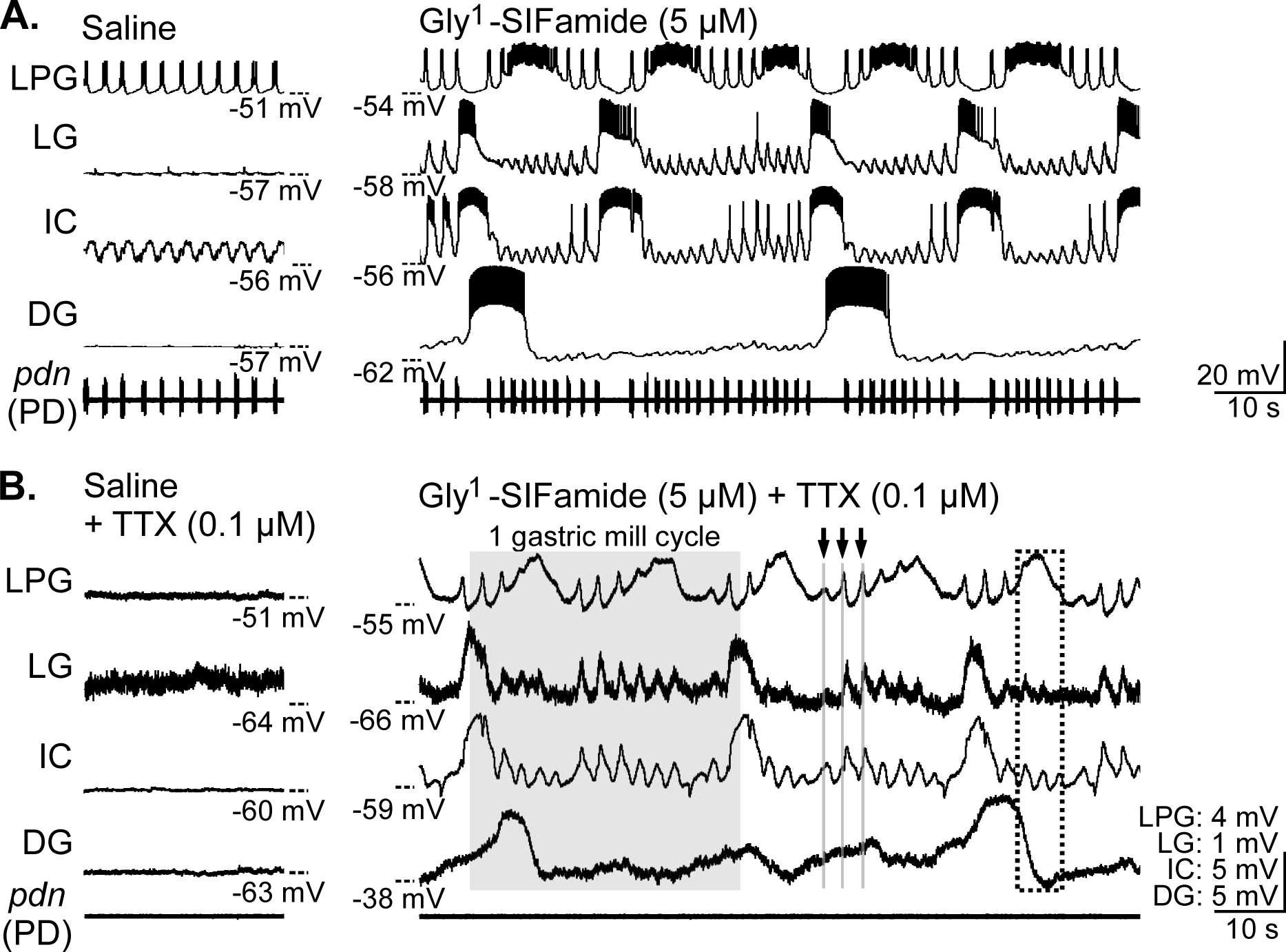
Graded synaptic transmission is sufficient for a coordinated gastric mill rhythm in Gly^1^-SIFamide. ***A***, Intracellular recordings of LPG, LG, IC, and DG during saline (***left***) and Gly^1^-SIFamide application (***right***). In saline, only the pyloric rhythm was active, with LPG bursting in pyloric time with the PD neuron (*pdn*). In Gly^1^-SIFamide, the gastric mill rhythm was activated (LG, DG, IC) and LPG generated both pyloric- and gastric mill-timed bursts. LPG pyloric-timed bursting in both saline and Gly^1^-SIFamide was time-locked to the PD neuron *(pdn*). ***B***, Tetrodotoxin (TTX) was applied to the preparation to block action potentials (***left***), then combined with Gly^1^-SIFamide (***right***). LPG, LG, IC, and DG generated gastric mill-timed oscillations in the absence of action potentials during TTX + Gly^1^-SIFamide application. The shaded region indicates one gastric mill cycle, using the LG neuron as a reference neuron. Note that LG and IC were co-active and out of phase with LPG and DG neurons, as in Gly^1^-SIFamide application without TTX (***A***). Troughs in DG oscillations appeared to alternate with peaks in LPG oscillations (*dotted box*). LPG, LG, and IC also generated pyloric-timed oscillations (arrows and grey lines), suggesting that Gly^1^-SIFamide also activated the pyloric rhythm without action potential-mediated transmission.

Our voltage-clamp analysis focused on whether there was modulation of bidirectional synapses between LPG and each of the other gastric mill neurons, as we were particularly interested in how well the switching neuron LPG is integrated into the gastric mill network in the Gly^1^-SIFamide modulatory condition. However, the triphasic pattern of the Gly^1^-SIFamide gastric mill rhythm requires chemical synapses from some or all of the LG, IC, and DG gastric mill neurons (21). Voltage-clamp analysis of all synaptic combinations is a daunting task and we instead turned to entrainment experiments to quantify the relative ability of each gastric mill neuron to regulate gastric mill-timed activity of each of the other gastric mill neurons in the intact network.

### Entrainment

To examine the ability of gastric mill neurons to regulate gastric mill bursting of the other neurons, we injected rhythmic depolarizing current into each entraining neuron (LG, IC, DG, or LPG [both copies]) across a range of burst frequencies (0.03 – 0.12 Hz; see Methods) and quantified its ability to entrain the burst frequency of the other neurons (Figs. 6 – 8; Supplemental Figs. S1 – S9).

**Figure 6.**
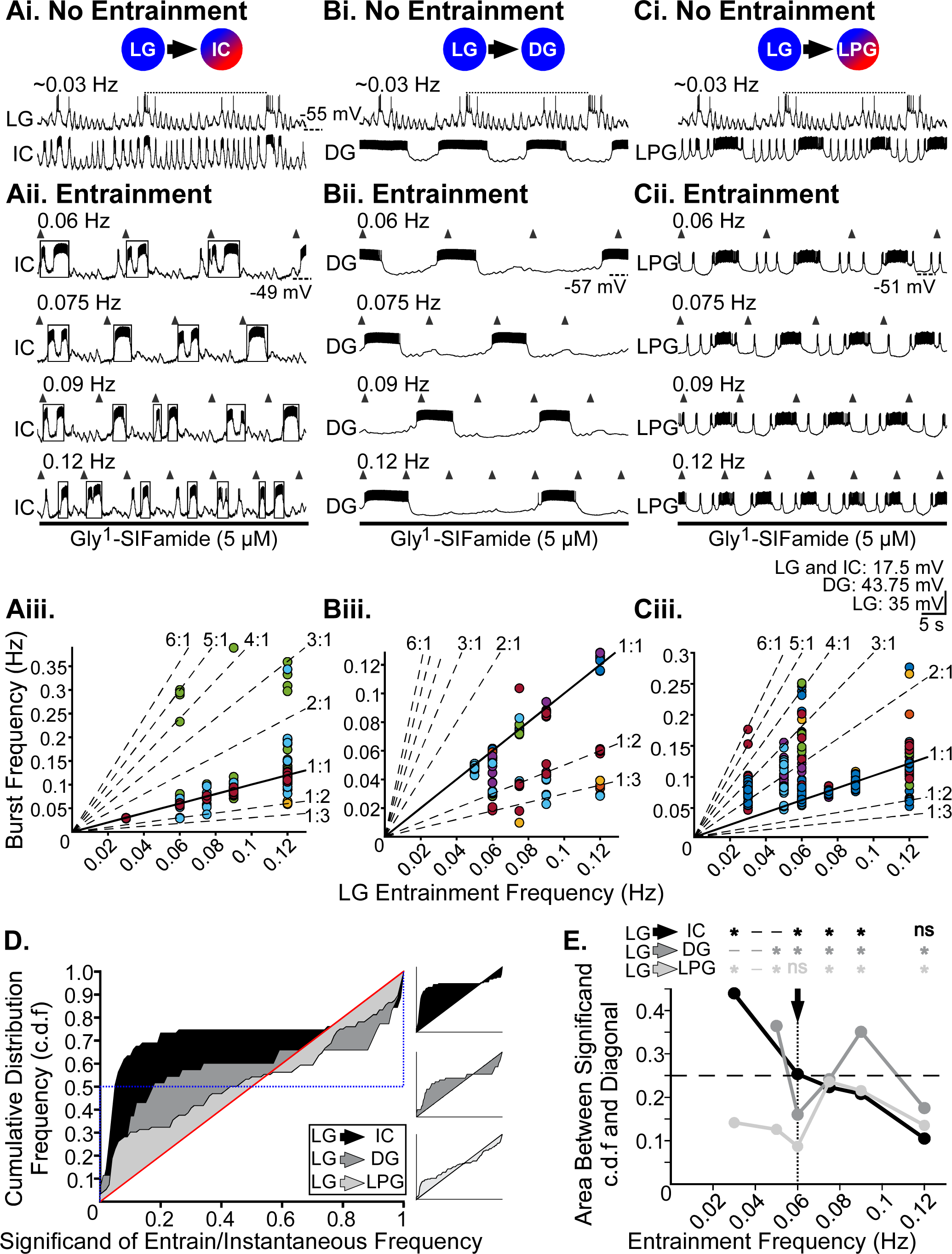
LG entrained IC, DG, and LPG gastric mill neurons during Gly^1^-SIFamide application. All columns are from the same preparation, in which LG entrainment of test neurons IC (***Ai-iii***), DG (***Bi-iii***), and LPG (***Ci-iii***) was examined. In Gly^1^-SIFamide (5 µM) prior to any manipulations, LG burst frequency was approximately 0.03 Hz and IC, DG, and LPG were bursting (***Ai-Ci***, No Entrainment). Dotted line indicates one LG cycle. ***Aii-Cii***, To determine the ability of LG to entrain the other neurons, LG was rhythmically depolarized (*upward arrowheads*, 5 s duration; LG recordings are omitted for clarity) at different entrainment frequencies, and IC (***Aii***), DG (***Bii***), and LPG (***Cii***) activity recorded. IC is active in both pyloric and gastric mill time during the Gly^1^-SIFamide rhythm, but only its gastric mill bursts were analyzed. IC gastric mill bursts (boxes) were identified as in previous studies (12, 21)(see Methods). ***Aiii-Ciii***, Instantaneous gastric mill burst frequency of IC (***Aiii***), DG (***Biii***), and LPG (***Ciii***) was plotted as a function of LG entrainment frequency. Each colored data point is the instantaneous burst frequency for one cycle (1/duration between two consecutive bursts), and each color represents data from a single preparation. Solid diagonal line indicates perfect entrainment (1:1), where instantaneous burst frequency was equal to entrainment frequency. Dashed diagonal lines indicate other modes of entrainment at which test neuron burst frequency would be a multiple, or a fraction, of entrainment frequency. Data points that fall between diagonal lines indicate a lack of LG entrainment of that neuron burst. Numbers of instantaneous burst frequencies plotted at each frequency are presented in Table 4. ***D***, The cumulative distribution frequency (c.d.f.) of significands are overlaid for LPG (*light grey*), DG (*dark grey*), and IC (*black*) responses to an LG entrainment frequency of 0.06 Hz. Significands are the value to the right of the decimal after dividing each instantaneous burst frequency of IC, LPG, or DG by the LG entrainment frequency. The red diagonal line represents a c.d.f. where significands are randomly distributed between 0 and 1, indicating no entrainment. The blue dotted line represents a c.d.f. with a large portion of the distribution close to 0 and/or 1, indicating entrainment. Shaded regions indicate the area between the c.d.f. for each neuron and the diagonal line. Insets of c.d.f. graphs enable visualization of area for each neuron separately. ***E***, Area between the significand c.d.f. curve and the diagonal line for IC (*black*), DG (*dark grey*), and LPG (*light grey*) as a function of LG entrainment frequency. The arrow and vertical dotted line indicate the c.d.f. area for the 0.06 Hz c.d.f. histograms plotted in (***D***) (other c.d.f.s are presented in Supplemental Fig. S1). The horizontal dashed line at 0.25 indicates one example of perfect entrainment, calculated as the area between the dotted blue line and the red diagonal line in (***D***). The c.d.f. for each neuron at each entrainment frequency was compared to a uniform distribution indicating zero integer coupling (i.e., diagonal line) to test for statistical significance using the Kolmogorov-Smirnoff test with a corrected alpha level of 0.007 (see Methods). Area values different from the uniform distribution at each entrainment frequency are indicated above the graph for each neuron pair. *p < 0.007, ^ns^p > 0.007 (see Table 5). Dash indicates data was not obtained at that entrainment frequency.

### LG entrainment of IC, DG, and LPG

We first examined whether LG could entrain IC, DG, and LPG neurons in Gly^1^-SIFamide (5 µM, Fig. 6). In an example experiment, prior to any manipulations, LG was bursting at approximately 0.03 Hz and IC, DG, and LPG were bursting in a relatively coordinated manner (Fig. 6Ai-Ci, “No Entrainment”). For clarity, for all entrainment frequencies presented, we use upward arrowheads to indicate the onset of each entrained neuron burst and only show recordings of the test neurons. When LG was stimulated with an entrainment frequency of 0.06 Hz (upward arrowheads), IC generated a slow burst with each LG stimulation (Fig. 6Aii, boxes). As LG entrainment frequency was increased, IC continued to generate 1-2 slow bursts within each LG stimulation cycle, with two slow IC bursts triggered by an LG stimulation more frequently at higher entrainment frequencies (Fig. 6Aii). IC bursts in both pyloric- and gastric mill-time. IC gastric mill bursts were identified as previously described (Fig. 6Aii, boxes; see Methods) (21). DG bursts overlapped with LG bursts in the No Entrainment condition, except for one LG cycle, where DG burst twice within the cycle (Fig. 6Bi, No Entrainment, dotted line). However, when the LG entrainment frequency was increased from 0.06 Hz to 0.09 Hz, DG generated one burst for every two LG cycles (Fig. 6Bii). At 0.12 Hz entrainment frequency, DG burst frequency slowed further, such that DG generated one burst for every four LG cycles (Fig. 6Bii). Thus, DG was coordinated with LG at higher entrainment frequencies but not in a 1:1 relationship. Finally, for LG and LPG slow bursting, without entrainment, LPG typically generated one burst per LG cycle, except for one cycle where LPG generated two slow bursts (Fig. 6Ci, No Entrainment). As the LG entrainment frequency was increased from 0.06 Hz to 0.12 Hz, LPG appeared to maintain 1:1 coordinated slow bursting (Fig. 6Cii). At 0.12 Hz LG entrainment frequency, a 1:1 relationship between LPG and LG persisted, although the relative timing between each LPG burst and the onset of the LG bursts was less consistent than at lower entrainment frequencies (Fig. 6Cii, 0.12Hz). LG does regulate LPG slow burst cycle period (21). The example experiment shown indicates that LG was able to entrain LPG within a range of frequencies (0.06 – 0.09 Hz), whereas above that, LG regulated the LPG cycle period but was not able to entrain LPG bursting.

Across preparations, to determine whether IC, DG, and LPG neurons were entrained to the LG entrainment frequencies, we plotted the instantaneous burst frequencies (1/time between two consecutive bursts) against LG entrainment frequency for all gastric mill-timed bursting, across all preparations (Fig. 6Aiii-Ciii). Each color represents one preparation. For LG and IC, the data points from each preparation clustered mostly on the solid diagonal line which indicates a 1:1 relationship of LG and IC bursting (Fig. 6Aiii). For instance, at 0.06 Hz, 0.075 Hz, and 0.09 Hz there are a total of 71, 48, and 71 cycles plotted, respectively. However, these data points are largely overlapping due to their entrainment by LG at these frequencies (Fig. 6Aiii; n = 3 – 7 preparations per frequency; see Table 4 for numbers of cycles at each frequency). At some frequencies, in some preparations, IC did not burst in a 1:1 manner, e.g., 0.06 Hz and 0.12 Hz (Fig. 6Aiii). Diagonal lines indicate other modes of entrainment in which the test neuron burst frequency was a multiple (or fraction) of the entrainment frequency (Fig. 6Aiii, dashed diagonal lines, see Methods). The addition of these lines enabled us to determine whether the data points not clustered on the 1:1 diagonal line were still entrained to the LG frequency by clustering at another entrainment mode, or perhaps not entrained if spread between entrainment modes. For instance, at 0.06 Hz, data points for instantaneous IC burst frequencies clustered primarily near the 1:1 line (65/71 data points). However, the additional points fell near the 1:2 (2/71), 4:1 (1/71), or 5:1 (3/71) lines, indicating that even if it was not a 1:1 relationship for every IC burst, IC activity was linked to that of LG and therefore was entrained by LG at 0.06 Hz. Conversely, at 0.12 Hz, while some data points fell on integer lines, many fell between them, indicating that IC was not readily entrained by LG at this frequency (Fig. 6Aiii).

**Table 4.** Number of test neuron cycles per preparation from each LG, IC, and DG entrainment frequency in Gly^1^-SIFamide.

| LG Entrainment Neuron |  |  |  | IC Entrainment Neuron |  |  |  | DG Entrainment Neuron |  |  |  |
| --- | --- | --- | --- | --- | --- | --- | --- | --- | --- | --- | --- |
| Test Neuron | Entrain Freq (Hz) | # Cycles / Prep | *n* | Test Neuron | Entrain Freq (Hz) | # Cycles / Prep | *n* | Test Neuron | Entrain Freq (Hz) | # Cycles / Prep | *n* |
| IC | 0.03 | 9-10 | 3 | LG | 0.03 | -- | - | LG | 0.03 | 9-18 | 3 |
|  | 0.04 | -- | - |  | 0.04 | 4-22 | 2 |  | 0.04 | 12-19 | 4 |
|  | 0.05 | -- | - |  | 0.05 | 8-13 | 4 |  | 0.05 | 7-18 | 6 |
|  | 0.06 | 8-14 | 7 |  | 0.06 | 6-11 | 5 |  | 0.06 | 5-11 | 8 |
|  | 0.075 | 8-10 | 6 |  | 0.075 | 2-9 | 6 |  | 0.075 | 3-10 | 8 |
|  | 0.09 | 10-11 | 7 |  | 0.09 | 5-8 | 4 |  | 0.09 | 2-9 | 8 |
|  | 0.12 | 4-14 | 7 |  | 0.12 | 4-6 | 4 |  | 0.12 | 1-9 | 7 |
| DG | 0.03 | -- | - | DG | 0.03 | 4-18 | 3 | IC | 0.03 | 12-14 | 2 |
|  | 0.04 | -- | - |  | 0.04 | 6-11 | 3 |  | 0.04 | 9-18 | 4 |
|  | 0.05 | 8-9 | 2 |  | 0.05 | 9-11 | 3 |  | 0.05 | 9-17 | 5 |
|  | 0.06 | 3-9 | 7 |  | 0.06 | 2-10 | 8 |  | 0.06 | 2-10 | 8 |
|  | 0.075 | 2-9 | 6 |  | 0.075 | 1-10 | 8 |  | 0.075 | 6-10 | 7 |
|  | 0.09 | 1-9 | 6 |  | 0.09 | 1-10 | 7 |  | 0.09 | 3-9 | 7 |
|  | 0.12 | 1-9 | 7 |  | 0.12 | 1-10 | 7 |  | 0.12 | 4-7 | 6 |
| LPG | 0.03 | 23-29 | 3 | LPG | 0.03 | 6-28 | 4 | LPG | 0.03 | 15-29 | 3 |
|  | 0.04 | -- | - |  | 0.04 | 15-19 | 3 |  | 0.04 | 10-21 | 5 |
|  | 0.05 | 14-19 | 2 |  | 0.05 | 6-12 | 4 |  | 0.05 | 6-18 | 6 |
|  | 0.06 | 9-20 | 8 |  | 0.06 | 7-10 | 9 |  | 0.06 | 9-16 | 11 |
|  | 0.075 | 8-10 | 7 |  | 0.075 | 3-10 | 9 |  | 0.075 | 7-14 | 10 |
|  | 0.09 | 8-10 | 8 |  | 0.09 | 3-9 | 8 |  | 0.09 | 5-12 | 11 |
|  | 0.12 | 5-9 | 8 |  | 0.12 | 1-11 | 8 |  | 0.12 | 4-10 | 11 |

For LG and DG, there was a range of instantaneous DG burst frequencies at each LG entrainment frequency (Fig. 6Biii; n = 2-7, Table 4). Unlike IC, DG was not locked to primarily one mode of entrainment. For example, in one preparation (Fig. 6Bii, light blue data points), DG was entrained 1:1 at 0.05 Hz, but at other entrainment frequencies DG exhibited a 1:2 (0.06 – 0.09 Hz) or 1:3 (0.09 – 0.12 Hz) relationship to LG bursting. Overall, LG to DG entrainment appeared to be weaker than LG to IC, e.g., note the spread of points between entrainment lines for an entrainment frequency of 0.06 Hz (Fig. 6Biii). However, it was a relatively small number of points (7/35 data points) that were spread between entrainment mode lines, the remainder of the data points clustered near 1:1 (12/35 data points), 1:2 (10/35), and 1:3 (6/35) lines. Thus, LG had some ability to entrain DG, although not always in a 1:1 relationship (Fig. 6Biii).

For LG and LPG slow bursting, we observed a range of instantaneous LPG slow burst frequencies at most entrainment frequencies, except at 0.075 and 0.09 Hz (Fig. 6Ciii, n = 2-8, Table 4). Although some data points fell on entrainment mode lines, many points, across multiple experiments, were spread between entrainment modes between 0.03 and 0.06 Hz, and for 0.12 Hz (Fig. 6Ciii). However, at 0.075 and 0.09 Hz, LPG data points clustered on the 1:1 entrainment mode, with little variability, suggesting that LG can entrain LPG within a limited frequency range (Fig. 6Ciii). While these plots enabled a qualitative assessment of entrainment, we pursued a quantitative analysis as well.

### Quantification of LG Entrainment of IC, DG, and LPG

To quantify entrainment between LG and IC, DG, and LPG, we collapsed the data points at each entrainment frequency into a single value using a previously described approach to quantify integer coupling relationships, e.g., 1:1, 2:1, etc., for entrainment neuron/test neuron burst frequencies (35). Specifically, we first calculated the ratio between each instantaneous neuron burst frequency and its respective LG entrainment frequency and then used the significands (value to the right of the decimal) from each set of ratios (i.e., IC/LG, DG/LG, LPG/LG) at each entrainment frequency to generate a cumulative probability distribution (c.d.f.) of significands (Fig. 6D; Supplemental Fig. S1). Perfect integer coupling would result in all significands being close to 0 or 1. However, if there was no relationship between entrainment frequency and test neuron burst frequency, then the significands would be distributed relatively evenly between 0 and 1, resulting in a c.d.f. close to a diagonal line (y = x) (Fig. 6D, red diagonal line) (35). For the example entrainment frequency of 0.06 Hz, c.d.f.s of significands (bin size 0.01) for each of the test neurons are overlaid (Fig. 6D; IC, black; DG, dark grey; LPG, light grey; remaining entrainment frequency c.d.f.s in Supplemental Fig. S1). The blue dotted line indicates an extreme in which 50% of a theoretical significand distribution is 0 and 50% is 1, indicating an example of perfect entrainment (Fig. 6D, blue dotted line). In the c.d.f. for IC significands (black) ∼70 % of significands were less than ∼0.1 (Fig. 6D, black shape). This shape reflects the clustering of IC instantaneous burst frequency around 1:1 coupling at 0.06 Hz entrainment in the scatter plot (Fig. 6Aiii). The c.d.f. of DG significands indicates that fewer were close to 0 or 1 (Fig. 6D, dark grey). This reflects the spread of some DG data points between entrainment modes at 0.06 Hz in the scatter plot (Fig. 6Biii). The c.d.f. of LPG significands indicated even weaker integer coupling at 0.06 Hz entrainment frequency, with the distribution being more similar to the diagonal line, demonstrating a relatively uniform distribution between 0 and 1 (Fig. 6D, light grey). This reflects the large spread of LPG data points between entrainment modes at 0.06 Hz (Fig. 6Ciii). Qualitatively, the c.d.f. histograms suggest that LG entrains IC more strongly than DG but does not entrain LPG at 0.06 Hz (Fig. 6D). Consistent with the scatter plots (Fig. 6Ciii), the c.d.f.s for LG to LPG entrainment at 0.075 and 0.09 indicate stronger integer coupling (Supplemental Fig. S1).

Once the c.d.f. for each test neuron, at each entrainment frequency (Fig. 6D; Supplemental Fig. S1), was determined, we calculated an overall measure of the extent of integer coupling at each entrainment frequency by calculating the area between each c.d.f. histogram (test neuron coupling) and the diagonal line (no integer coupling) (Fig. 6D, shaded regions). These area measurements for each neuron were plotted against entrainment frequency, including from the example c.d.f.s for 0.06 Hz in figure 6D (Fig. 6E, downward arrow/dotted line). The dashed line at y = 0.25 indicates an example of perfect integer coupling, where test neuron burst frequency is equal to the entrainment neuron frequency, i.e., the area between the blue dotted line and the red diagonal line in figure 6D (35). To determine whether the c.d.f. for each neuron at each entrainment frequency was significantly different from a uniform distribution (i.e., the diagonal line), we used a two-sample Kolmogorov-Smirnov test to compare each c.d.f. to a uniform distribution. A corrected significance level of 0.007 (see Methods) for the comparison between test neuron c.d.f. and a uniform distribution was used as the criterion for entrainment at a particular frequency. Results of this statistical analysis is indicated above each entrainment frequency for each neuron (Fig. 6E). For instance, the area between the IC c.d.f. and the diagonal line at 0.03 Hz entrainment frequency was approximately 0.45 but the IC c.d.f. area decreased at higher entrainment frequencies (Fig. 6E, black line/symbols). Specifically, LG-to-IC integer coupling, and thus entrainment, occurred between 0.03 – 0.09 Hz, but not at 0.12 Hz (Fig. 6E; Table 5). Although the area between c.d.f. and the diagonal line for DG was greater at an entrainment frequency of 0.05 and 0.075 Hz, than at other frequencies (Fig. 6E, dark grey line/circles), DG was entrained by LG at all entrainment frequencies (Table 5). Finally, the area for LPG was below 0.25 across frequencies but was non-zero (Fig. 6E, light grey line/circles) and this area measurement was different from the uniform distribution at all entrainment frequencies except 0.06 Hz (Table 5). Overall, these results indicate that IC, DG, and LPG were integer coupled to and entrained by LG to some extent, supporting our voltage-clamp analysis that there was a functional synapse between these neurons in Gly^1^-SIFamide.

**Table 5.** Statistical analysis of the ability of LG, IC, and DG to entrain the other gastric mill neurons during Gly^1^-SIFamide application.

| Entrainment Frequency | p value (D) |  |  |  |  |  |  |  |  |
| --- | --- | --- | --- | --- | --- | --- | --- | --- | --- |
|  | Entrainment/Test Neuron |  |  |  |  |  |  |  |  |
|  | LG/IC | LG/DG | LG/LPG | IC/LG | IC/DG | IC/LPG | DG/LG | DG/IC | DG/LPG |
| 0.03 | <0.001<br>(0.902) | --- | <0.001<br>(0.284) | --- | 0.271<br>(0.137) | <0.001<br>(0.324) | <0.001<br>(0.353) | 0.003<br>(0.245) | 0.025<br>(0.206) |
| 0.04 | --- | --- | --- | 0.001<br>(0.265) | <0.001<br>(0.382) | <0.001<br>(0.275) | 0.360<br>(0.127) | 0.074<br>(0.176) | <0.001<br>(0.324) |
| 0.05 | --- | <0.001<br>(0.725) | 0.001<br>(0.265) | 0.051<br>(0.186) | <0.001<br>(0.520) | <0.001<br>(0.363) | <0.001<br>(0.353) | <0.001<br>(0.373) | <0.001<br>(0.441) |
| 0.06 | <0.001<br>(0.509) | <0.001<br>(0.324) | 0.034<br>(0.196) | 0.0057<br>(0.235) | <0.001<br>(0.275) | <0.001<br>(0.451) | <0.001<br>(0.600) | <0.001<br>(0.647) | 0.002<br>(0.255) |
| 0.075 | <0.001<br>(0.520) | <0.001<br>(0.559) | <0.001<br>(0.520) | 0.051<br>(0.186) | <0.001<br>(0.461) | <0.001<br>(0.608) | <0.001<br>(0.627) | <0.001<br>(0.529) | 0.014<br>(0.216) |
| 0.09 | <0.001<br>(0.431) | <0.001<br>(0.676) | <0.001<br>(0.461) | 0.034<br>(0.196) | <0.001<br>(0.539) | <0.001<br>(0.422) | <0.001<br>(0.412) | <0.001<br>(0.392) | 0.001<br>(0.265) |
| 0.12 | 0.074<br>(0.176) | <0.001<br>(0.392) | <0.001<br>(0.284) | <0.001<br>(0.353) | <0.001<br>(0.520) | 0.003<br>(0.245) | <0.001<br>(0.471) | <0.001<br>(0.363) | 0.051<br>(0.186) |

Two-sample Kolmogorov-Smirnov test was used to compare each c.d.f. to a uniform distribution, alpha level was 0.007 to correct for multiple comparisons

### Entrainment and integer coupling among the other gastric mill neurons

For the remainder of the neurons, we focused on the quantification afforded by generating c.d.f.s of significands at each entrainment frequency, and measuring the area between the test neuron c.d.f. and a uniform distribution (no coupling). The complete assessment of the ability of IC, DG, and LPG to entrain the other neurons, including individual data points as in figure 6 are provided in Supplemental Figs. S2-S9. Overall, we found that all neurons (LG, IC, DG, LPG) could entrain all other neurons, at least at some entrainment frequencies, however the extent of entrainment was variable (Figs. 6 – 8, Supplemental Figs. S1 - S9).

At an example IC entrainment frequency of 0.06 Hz (additional c.d.f.s in Supplemental Fig. S3), IC most effectively entrained LPG (Fig. 7Ai, black) with ∼60% of LPG/IC significands being less than 0.1 (Fig. 7Ai; n = 3-9, Table 4). The c.d.f.s for DG (dark grey) and LG (light grey) were closer to the diagonal line, with ∼25% and ∼20% of significands less than 0.1, respectively (Fig. 7Ai). The c.d.f. of significands for IC/LG (Fig. 7Ai, light grey) was fairly close to the uniform distribution (red diagonal line). In fact, using the c.d.f.s compared to a uniform distribution, IC entrained LG at 0.04, 0.06, and 0.12 Hz, but not at 0.05, 0.075, or 0.09 Hz (Fig. 7Aii; Table 5). LG appeared better able to entrain IC as the LG/IC area measurements were farther away from zero (compare Fig. 6D and Fig. 7Aii). IC entrained DG from 0.04 – 0.12 Hz, but not at 0.03 Hz (Fig. 7Aii, dark grey circles and line; Table 5). Finally, LPG was entrained by IC at all entrainment frequencies (Fig. 7Aii; Table 5; Supplemental Figs. S2-S3). Thus, LG, DG, and LPG were all integer coupled to and entrained by IC at some entrainment frequencies, also supporting the presence of functional synapses from IC to LG, DG, and LPG in Gly^1^-SIFamide.

**Figure 7.**
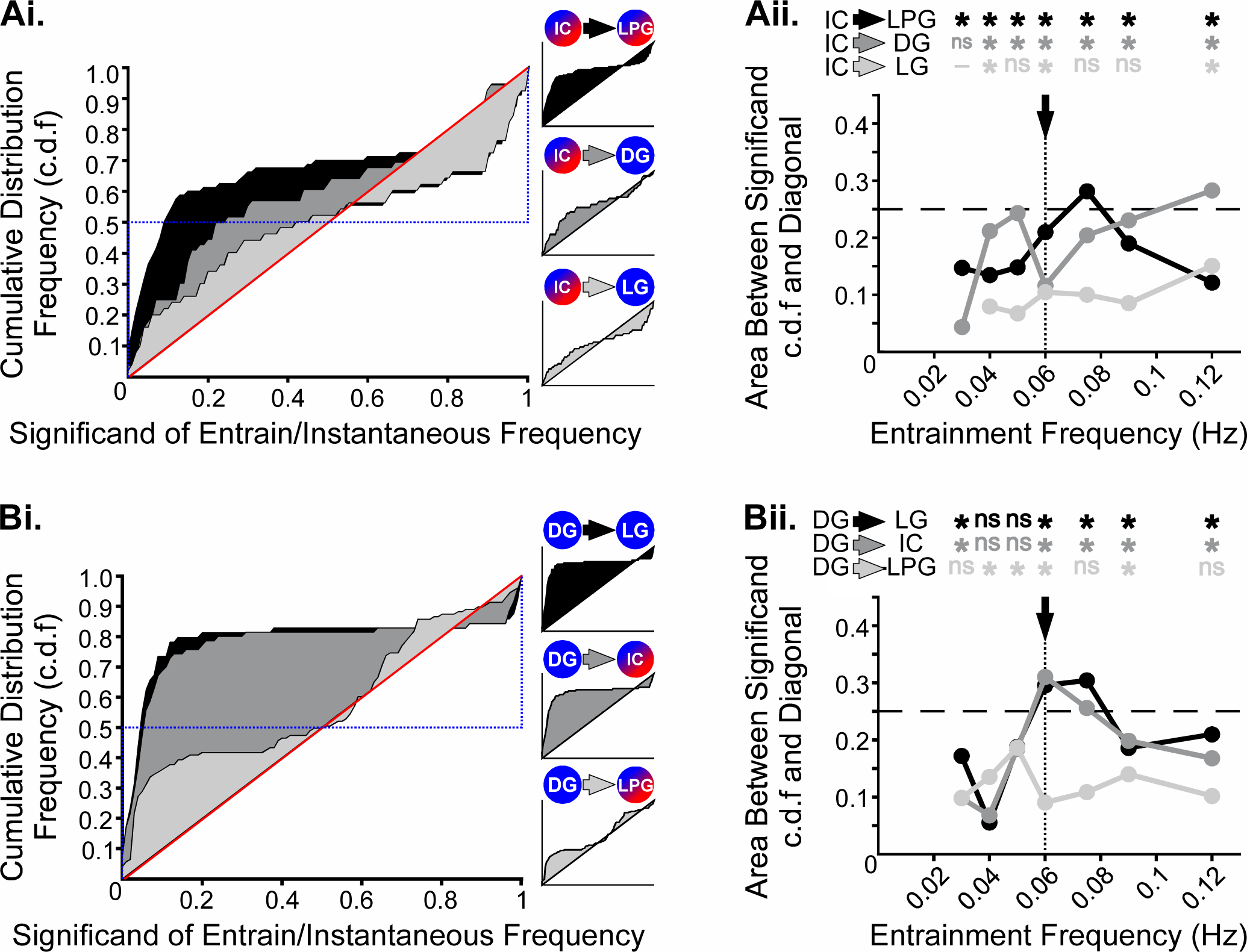
IC and DG entrained the other gastric mill neurons during Gly^1^-SIFamide application at some entrainment frequencies. ***Ai***, Overlaid cumulative distribution frequency histograms (c.d.f.s) for IC entrainment frequency of 0.06 Hz for LG (*light grey*), DG (*dark grey*), and LPG (*black*). ***Bi***, Overlaid c.d.f.s for DG entrainment frequency of 0.06 Hz for LPG (*light grey*), IC (*dark grey*), and LG (*black*). Area between the significand c.d.f. curve and the red diagonal line for LG (*light grey*), DG (*dark grey*), and LPG (*black*) as a function of IC entrainment frequency (***Aii***) or LPG (*light grey*), IC (*dark grey*), and LG (*black*) as a function of DG entrainment frequency (***Bii***). The arrow and vertical dotted lines indicate the c.d.f. area calculated for the 0.06 Hz c.d.f. histograms plotted in (***Ai, Bi***) (other c.d.f.s are presented in Supplemental Figs. S3, S5). The c.d.f. for each neuron at each entrainment frequency was compared to a uniform distribution indicating zero integer coupling (i.e., diagonal line) to test for statistical significance using the Kolmogorov-Smirnoff test with a corrected alpha level of 0.007 (See Methods). Area values different from the uniform distribution at each entrainment frequency are indicated above the graph for each neuron pair. *p < 0.007, ^ns^p > 0.007 (see Table 5). Dash indicates data was not obtained at that entrainment frequency.

The DG neuron is the most likely to be uncoordinated with the other gastric mill neurons in the Gly^1^-SIFamide gastric mill rhythm (21, 28). Despite this, stimulating DG across a range of entrainment frequencies revealed some degree of control over the burst frequency of the other neurons. For example, DG entrainment of LG and IC at an entrainment frequency of 0.06 Hz, was evident in the ∼75-80% of significands that were less than 0.1 for LG and IC, while there were ∼35% of significands less than 0.1 for LPG (Fig. 7Bi, shaded regions, LG (black); IC (dark grey); LPG (light grey) (additional c.d.f.s in Supplemental Fig. S5). The area between each c.d.f. and the diagonal line indicated that integer coupling was variable for DG to LG (black), IC (dark grey), and LPG (light grey) (Fig. 7Bii). LG and IC were entrained to DG at 0.03 and 0.05 – 0.12 Hz, but not entrained at 0.04 Hz (Fig. 7Bii; Table 5). LPG was entrained by DG at 0.04 – 0.06 and 0.09 Hz, but not entrained at 0.03, 0.075, or 0.12 Hz entrainment frequencies (Fig. 7Bii, Table 5; Supplemental Figs. S4-S5). The entrainment results correlate with the voltage-clamp analysis in that the DG to LPG synapse was weaker and more variable than LG or IC to LPG synapses (Figs. 2-4).

In the Gly^1^-SIFamide modulatory state, LPG more strongly entrained the DG neuron compared to IC and LG. In the c.d.f. histogram for DG at a 0.06 Hz entrainment frequency, ∼45% of significands were less than 0.1, and the area between DG c.d.f. and the diagonal was 0.21 (Fig. 8Ai-ii, black shaded region and black data point at downward arrow, respectively). Based on calculations of area between the c.d.f.s and the diagonal line of no entrainment (Fig. 8Ai-ii; Supplemental Figs. S6-S7), LG was entrained by LPG at 0.03, 0.075, and 0.09 Hz, but not at 0.04 – 0.06 or 0.12 Hz (Fig. 8Aii; Table 6). DG was entrained by LPG at all entrainment frequencies (Fig. 8Aii; Table 6) and IC was entrained by LPG only at 0.075 and 0.09 Hz, and not at the other frequencies (Fig. 8Aii; Table 6; Supplemental Figs. S6-S7).

**Figure 8.**
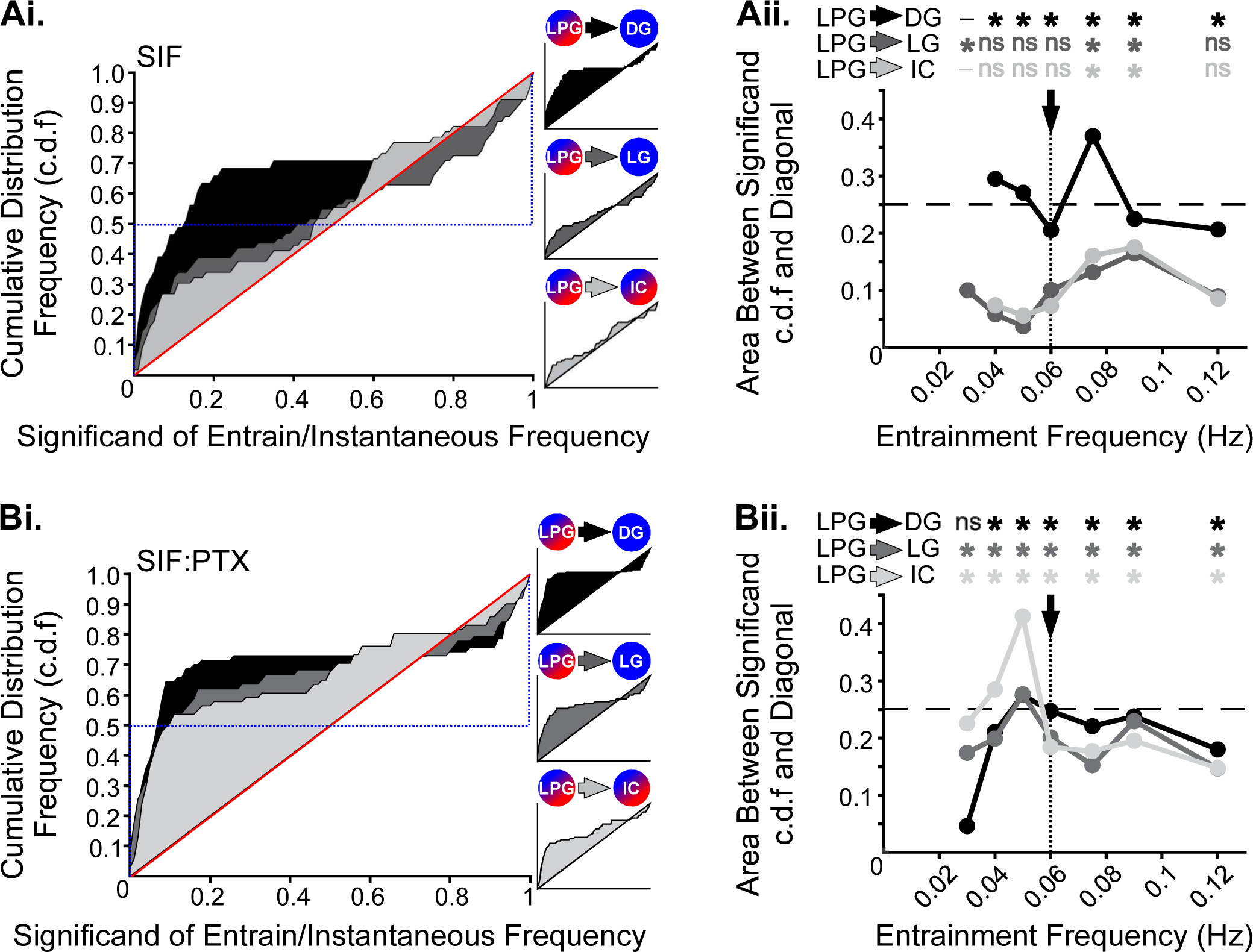
LPG entrainment of LG, IC, and DG gastric mill neurons during Gly^1^-SIFamide (SIF) and Gly^1^-SIFamide plus picrotoxin (SIF:PTX) application. ***Ai***, Overlaid cumulative distribution frequency histograms (c.d.f.s) for LPG entrainment frequency of 0.06 Hz for IC (*light grey*), LG (*dark grey*), and DG (*black*) during Gly^1^-SIFamide (SIF) application. ***Bi***, Overlaid c.d.f.s for LPG entrainment frequency of 0.06 Hz for IC (*light grey*), LG (*dark grey*), and DG (*black*) during Gly^1^-SIFamide plus picrotoxin (SIF:PTX) application. Area between the significand c.d.f. curve and the red diagonal line for IC (*light grey*), LG (*dark grey*), and DG (*black*) as a function of LPG entrainment frequency during SIF (***Aii***) or SIF:PTX (***Bii***). The arrow and vertical dotted lines indicate the c.d.f. area calculated for the 0.06 Hz c.d.f. histograms plotted in (***Ai, Bi***) (other c.d.f.s are presented in Supplemental Figs. S7, S9). The c.d.f. for each neuron at each entrainment frequency was compared to a uniform distribution indicating zero integer coupling (i.e., diagonal line) to test for statistical significance using the Kolmogorov-Smirnoff test with a corrected alpha level of 0.007 (See Methods). Area values different from the uniform distribution at each entrainment frequency are indicated above the graph for each neuron pair. *p < 0.007, ^ns^p > 0.007 (see Table 6). Dash indicates data was not obtained at that entrainment frequency.

**Table 6.** Number of test neuron cycles per preparation from each LPG entrainment frequency in Gly^1^-SIFamide (SIF) and Gly^1^-SIFamide plus PTX (SIF:PTX), and statistical analysis of entrainment.

| LPG Entrainment Neuron<br>(SIF) |  |  |  |  | LPG Entrainment Neuron<br>(SIF:PTX) |  |  |  |  |
| --- | --- | --- | --- | --- | --- | --- | --- | --- | --- |
| Test<br>Neuron | Entrain<br>Freq<br>(Hz) | #<br>Cycles/<br>Prep | *n* | p value (D) | Test<br>Neuron | Entrain<br>Freq<br>(Hz) | #<br>Cycles/<br>Prep | *n* | p value (D) |
| LG | 0.03 | 10-20 | 2 | <0.001 (0.314) | LG | 0.03 | 11-21 | 4 | <0.001 (0.363) |
|  | 0.04 | 10-17 | 4 | 0.271 (0.137) |  | 0.04 | 8-27 | 4 | <0.001 (0.353) |
|  | 0.05 | 9-18 | 5 | 0.689 (0.098) |  | 0.05 | 9 | 3 | <0.001 (0.549) |
|  | 0.06 | 5-16 | 6 | 0.009 (0.225) |  | 0.06 | 7-12 | 7 | <0.001 (0.441) |
|  | 0.075 | 5-12 | 6 | <0.001 (0.304) |  | 0.075 | 4-9 | 7 | <0.001 (0.333) |
|  | 0.09 | 4-9 | 5 | <0.001 (0.333) |  | 0.09 | 1-9 | 7 | <0.001 (0.520) |
|  | 0.12 | 3-10 | 5 | 0.015 (0.216) |  | 0.12 | 1-9 | 7 | <0.001 (0.275) |
| IC | 0.03 | -- | - | -- | IC | 0.03 | 7-14 | 4 | <0.001 (0.431) |
|  | 0.04 | 10-12 | 4 | 0.034 (0.196) |  | 0.04 | 1-11 | 5 | <0.001 (0.618) |
|  | 0.05 | 9-17 | 5 | 0.020 (0.147) |  | 0.05 | 6-10 | 4 | <0.001 (0.814) |
|  | 0.06 | 5-16 | 5 | 0.034 (0.196) |  | 0.06 | 5-16 | 8 | <0.001 (0.412) |
|  | 0.075 | 4-12 | 5 | <0.001 (0.373) |  | 0.075 | 1-10 | 8 | <0.001 (0.373) |
|  | 0.09 | 3-9 | 5 | <0.001 (0.314) |  | 0.09 | 3-12 | 8 | <0.001 (0.402) |
|  | 0.12 | 3-9 | 5 | 0.051 (0.186) |  | 0.12 | 2-8 | 8 | <0.001 (0.353) |
| DG | 0.03 | -- | - | -- | DG | 0.03 | 11-16 | 4 | 0.356 (0.127) |
|  | 0.04 | 6-10 | 4 | <0.001 (0.637) |  | 0.04 | 9-12 | 5 | <0.001 (0.461) |
|  | 0.05 | 7-12 | 5 | <0.001 (0.559) |  | 0.05 | 9-10 | 4 | <0.001 (0.569) |
|  | 0.06 | 1-10 | 7 | <0.001 (0.451) |  | 0.06 | 5-10 | 8 | <0.001 (0.539) |
|  | 0.075 | 2-10 | 7 | <0.001 (0.696) |  | 0.075 | 3-9 | 8 | <0.001 (0.5) |
|  | 0.09 | 2-10 | 7 | <0.001 (0.471) |  | 0.09 | 3-10 | 8 | <0.001 (0.490) |
|  | 0.12 | 1-10 | 7 | <0.001 (0.480) |  | 0.12 | 2-8 | 7 | <0.001 (0.392) |

Two-sample Kolmogorov-Smirnov test to compare each c.d.f. to a uniform distribution, alpha level was 0.007 to correct for multiple comparisons

LPG can coordinate IC, LG, and DG during Gly^1^-SIFamide application when all other synapses are blocked. However, LPG was not necessary for coordinating these neurons when other chemical synapses among the gastric mill neurons were intact (21), suggesting overlapping functions of synaptic connections, which might interfere with measurements of the ability of LPG to entrain the other neurons. Thus, we performed an additional set of experiments to assess the ability of LPG to entrain LG, IC, and DG neurons in the presence of Gly^1^-SIFamide (SIF), plus picrotoxin (PTX, 10^−5^ M; Fig. 8Bi-ii). As noted above, PTX application blocks glutamatergic inhibitory synapses, isolating LPG from the glutamatergic LG, IC, and DG gastric mill neurons, but LPG cholinergic transmission is not blocked by PTX (21, 30, 32, 50). Test neuron c.d.f.s in SIF:PTX (Fig. 8Bi) appeared larger than in Gly^1^-SIFamide only (Fig. 8Ai). In SIF:PTX the c.d.f. histograms for LG (dark grey), IC (light grey), and DG (black) had ∼50-70% of significands that were less than 0.1 at 0.06 Hz entrainment frequency (Fig. 8Bi, shaded regions). Across all entrainment frequencies, most c.d.f.s appeared to indicate greater entrainment in SIF:PTX compared to Gly^1^-SIFamide alone (Supplemental Figs. S7 vs. S9). Quantitatively, with glutamatergic synapses blocked (SIF:PTX), LPG entrained DG between 0.04 – 0.12 Hz, but not at 0.03 Hz (Fig. 8Bii; Table 6). LPG was already able to entrain DG from 0.04 – 0.12 Hz in Gly^1^-SIFamide (Fig. 8Aii), but data was not obtained at 0.03 Hz in this condition. However, for LG and IC, which were entrained by LPG from 0.04 – 0.12 Hz in Gly^1^-SIFamide (Fig. 8Aii), in SIF:PTX LPG entrained them at all frequencies (Fig. 8Bii, LG, dark grey line/circles; IC, light grey line/circles) (Fig. 8Bii; Table 6; Supplemental Figs. S8-S9). Thus, it appears likely that synaptic actions from LG, IC, and/or DG in Gly^1^-SIFamide interfere with the ability of LPG to control LG and IC bursting. Because only cholinergic LPG synapses were active in SIF:PTX, our entrainment assay indicates direct functional synapses from LPG onto LG, IC, and DG neurons in the Gly^1^-SIFamide modulatory state, in alignment with our voltage-clamp findings (Figs. 2-4).

## Discussion

In this study we determined that there are functional, bidirectional synapses between LPG and gastric mill network neurons LG, IC, and DG in the modulatory state elicited by the neuropeptide Gly^1^-SIFamide. As revealed by two electrode voltage-clamp recordings, most of the synaptic connections are ineffective in saline conditions, but are enhanced by Gly^1^-SIFamide (Fig. 9). Additionally, we found that each neuron, including the switching neuron LPG, is capable of entraining other neurons in the network, within a physiological frequency range during the Gly^1^-SIFamide modulatory state. Thus, in addition to modulating intrinsic properties of LPG, thereby enabling dual frequency oscillations (21, 22), the neuropeptide Gly^1^-SIFamide enables previously ineffective synapses to become a functional pathway for a switching neuron to both be influenced by, and to influence, neurons in a network to which it transiently belongs.

**Figure 9.**
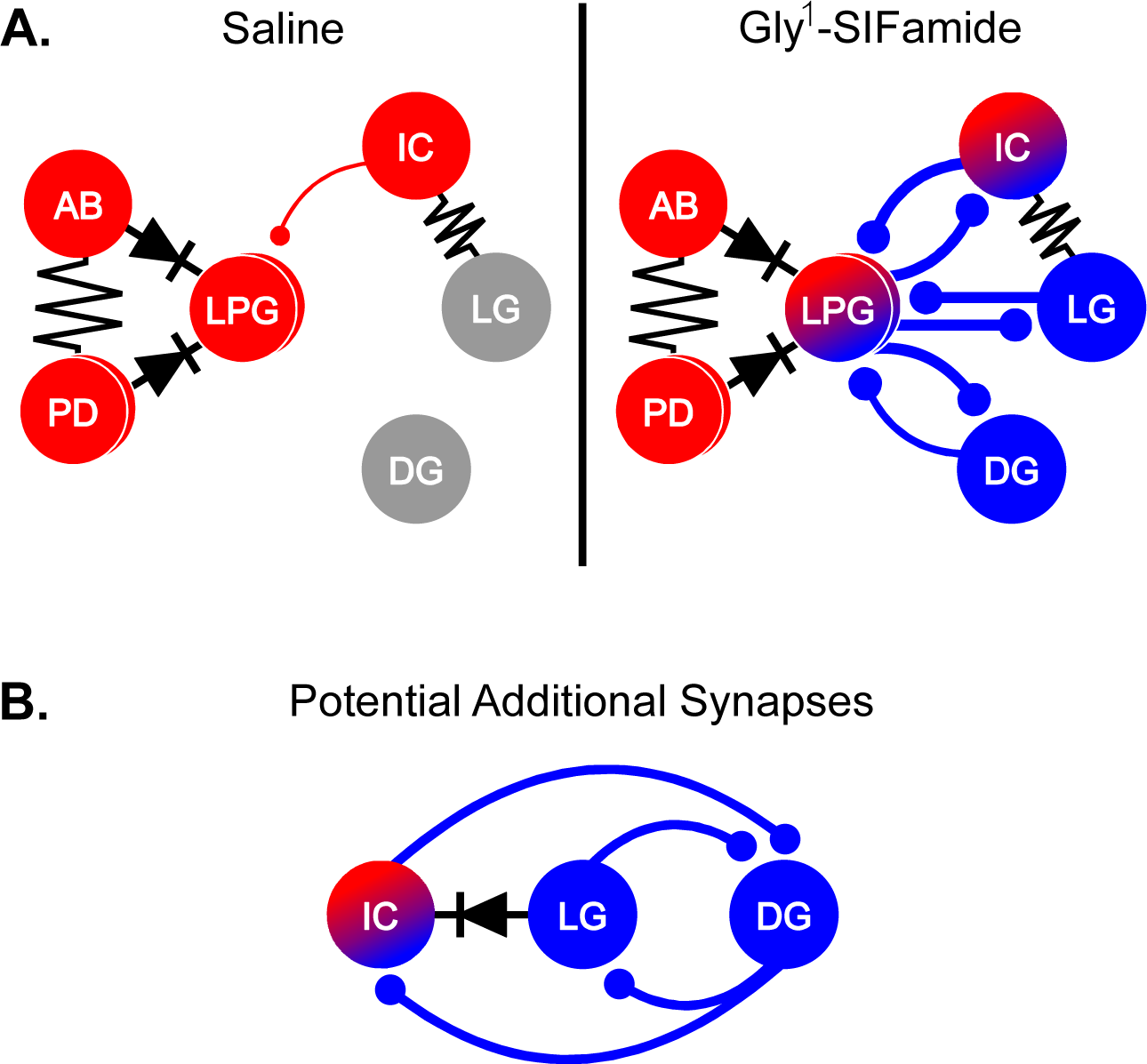
Gly^1^-SIFamide modulates multiple synapses. Updated partial circuit diagram from figure 1. ***A****, **Left**,* In saline conditions (Saline), the pyloric neuron LPG is electrically coupled to the AB/PD pyloric pacemaker ensemble, and receives a weak chemical inhibitory synapse from IC. The gastric mill rhythm is off. ***Right***, During Gly^1^-SIFamide application, LPG intrinsic properties are modulated to enable LPG switching into dual pyloric plus gastric mill activity (red/blue LPG circle). Gly^1^-SIFamide modulation also enhances bidirectional chemical inhibitory synapses between LPG and the IC, LG, and DG gastric mill neurons. ***B***, Potential additional inhibitory chemical synapses that are functional during Gly^1^-SIFamide application among the LG, IC, and DG neurons, based on results of entrainment experiments. Additionally, entrainment data suggests that the established electrical coupling between IC and LG may be rectifying (*diode*), such that positive current flows better from LG to IC. Resistor and diode, electrical coupling; ball and stick, chemical inhibition, thickness indicates relative synapse strength; red circles, pyloric neurons; blue circles, gastric mill neurons; red/blue circles; gastropyloric neurons.

### Modulation of Internetwork Synapses

Neuromodulator regulation of synaptic strength contributes to CPG flexibility (39, 51). In particular, neuropeptide modulation of synapses occurs throughout nervous systems with both pre- and postsynaptic sites of action (52–57). Modulator-elicited enhancement of internetwork synapses can serve as the sole mechanism by which a neuron is both recruited to generate activity at a second frequency, and coordinated with other neurons (6, 8, 20). In the saline state, LPG activity is pyloric due to electrical coupling with the pyloric pacemaker ensemble (12, 58). Gly^1^-SIFamide modulates multiple ionic currents to switch LPG into dual pyloric and gastric mill-timed bursting (12, 22). However, we found that in saline, among gastric mill neurons and LPG, there was only a weak synapse from IC onto LPG. Thus, modulation of intrinsic properties is not sufficient to coordinate LPG with other gastric mill neurons. The ability of Gly^1^-SIFamide to also enhance inhibitory connections between LPG and gastric mill neurons LG, IC, and DG, explain the ability of LPG and LG, IC, and DG to entrain each other (this study) and to generate a coordinated rhythm (21, 28), despite being functionally disconnected in saline conditions (Fig. 9A). Thus, parallel modulatory actions impact a switching neuron such that (1) modulation of intrinsic properties enables generation of rhythmic activity at a second frequency, and (2) modulation of synaptic properties between switching and network neurons enables coordination within the second network.

Regardless of whether synaptic modulation occurs in isolation or with modulation of intrinsic properties, network output is determined by complex interactions between both sets of properties (59). Such interactions may explain the diversity of entrainment we observed. Most simply, entrainment is 1:1 coordination between two neurons or rhythmic patterns, with test neuron frequency equal to entrainment frequency (60–63). While 1:1 entrainment among LPG, LG, IC, and DG occurred (notably from LG to IC), we also observed other coordination patterns. For instance, regardless of the entrainment neuron, DG exhibited some 1:2 entrainment (one DG burst for every two entrainment neuron bursts). In addition to LPG, DG is an endogenous burster in Gly^1^-SIFamide (21). Slower DG bursting at higher entrainment frequencies suggests that rhythmic synaptic inhibition may not be able to overcome the slower ionic currents underlying endogenous DG bursting, reinforcing the combined contributions of synaptic and intrinsic properties to network output.

Although communication was bidirectional between all pairs, differences in synaptic properties were observed. For example, LPG cholinergic output synapses onto the gastric mill neurons have a slower time course than glutamatergic synapses onto LPG, as do cholinergic versus glutamatergic synapses within the pyloric network (47, 48). Pharmacological block of glutamatergic transmission makes it likely that the LPG to DG synapse is direct. However, determining direct connections is difficult, even in small networks due to the complicating factor of highly prevalent electrical coupling, common across networks (64–66). For example, LPG-elicited synaptic currents in LG and IC are difficult to interpret due to LG/IC electrical coupling. Identifying direct synapses among gastric mill neurons would require photoinactivation to selectively eliminate subsets of neurons (47).

In addition to chemical synapses, properties of electrical synapses among gastric mill neurons are likely important in the Gly^1^-SIFamide rhythm. During Gly^1^-SIFamide, LG/IC gastric mill-timed bursting is typically coincident (21), likely due to electrical coupling (64). However, while LG entrained IC in a mostly 1:1 fashion across most frequencies, IC only entrained LG at half of the frequencies. We speculate this difference is due to either rectifying electrical coupling, where positive current flows better from LG to IC, or an inhibitory synapse from IC to LG (67) that is stronger than LG/IC electrical coupling. Rectification promotes CPG flexibility by enabling differential modulatory actions on coupled neurons, and may be important for neuronal switching between networks (11, 68–70). For instance, rectifying LPG/PD electrical coupling (58) may be important for enabling LPG switching into gastric mill timing (S-RH Fahoum, F Nadim, and DM Blitz, unpublished observations). Similarly, LG/IC rectification could enable IC recruitment from pyloric-only to dual pyloric/gastric mill timing, while preventing LG from being pulled into pyloric-timed activity. Thus, rectification can play an important role in maintaining network distinctions, even as some neurons are recruited to participate in secondary networks.

### Functional Implications of Synaptic Modulation

Synaptic modulation is a common mechanism for regulating the relative timing of neurons, and thus the particular activity pattern, although modulation of ionic currents that alter responses to synaptic input can also be important for patterning network output (1, 59, 71, 72). Other gastric mill rhythm versions are biphasic (73–75), whereas the Gly^1^-SIFamide-elicited rhythm has a third phase, during which endogenous LPG slow bursting is out of phase with the other neurons ((21). Inhibitory synapses from LPG to LG, IC, and DG likely ensure the offset between LPG and the other neurons, while synapses between LG, IC, and DG (Fig. 9B) are likely important for maintaining offsets in their timing. In fact, the rhythm collapses into two phases (LG/IC/DG co-active and alternating with LPG) when glutamatergic synapses are blocked via PTX, leaving only cholinergic LPG synapses functional (21). Additionally, LPG regulates LG and DG neuron firing frequency in Gly^1^-SIFamide (21). Collectively, these results emphasize the full incorporation of an endogenously bursting switching neuron, such that it can contribute to pattern generation in a second network.

Spike-mediated and graded neurotransmission play complementary roles at vertebrate and invertebrate synapses and are both subject to neuromodulation (55, 76–81). In the STNS, spike-mediated and graded transmission occur, but graded transmission is dominant and can be sufficient to generate fully coordinated pyloric rhythms under some modulatory states (43, 44, 46). Here we determined that a coordinated gastric mill rhythm can occur without spike-mediated transmission in the Gly^1^-SIFamide state. However, whether rhythm generating mechanisms differ for Gly^1^-SIFamide compared to other gastric-mill rhythm activators, and whether they require spike-mediated transmission is unknown (75, 82). Rhythm generation of episodic behaviors (e.g., chewing, locomotion) typically relies on intra-network synapses, as opposed to pacemaker neurons generating continuously active rhythms (e.g., pyloric, respiratory) (1, 83). In other gastric mill rhythm versions, reciprocal inhibition between the LG and Int1 neurons forms the core rhythm generator. Gly^1^-SIFamide elicits endogenous bursting in LPG and DG, but IC and LG cannot generate rhythmic activity in the absence of chemical synapses in Gly^1^-SIFamide (12, 21, 22). Thus, in addition to coordinating endogenous oscillations of the LPG switching neuron plus DG with other network neurons, the modulated graded synapses are likely necessary for rhythmic activity in at least some of the network neurons.

## Conclusions

There is increasing evidence from small- to large-scale, and local to long-distance, that neuromodulators can rapidly alter network composition based on sensory information, a required behavior, or required coordination between multiple behaviors (14–17, 19, 84–86). Our results indicate that parallel modulation of intrinsic and synaptic properties of a switching neuron can mediate recruiting a neuron into generating rhythmic activity at a second frequency (intrinsic properties) and coordinating those oscillations with other neurons in the network (synaptic properties), potentially providing additional flexibility under different behavioral states. It is also possible that the combined influence of parallel actions is necessary based on the particular complement of intrinsic and synaptic properties that must be overcome for a switching neuron to “escape” from its home network.

## SUPPLEMENTAL MATERIAL

Supplemental Figs. S1–S9: https://figshare.com/articles/figure/Supplemental_Synapses_pdf/25559007

## Acknowledgements

This work supported by the National Science Foundation IOS:1755283 (DMB).

---

**Supplemental Figure S1.**
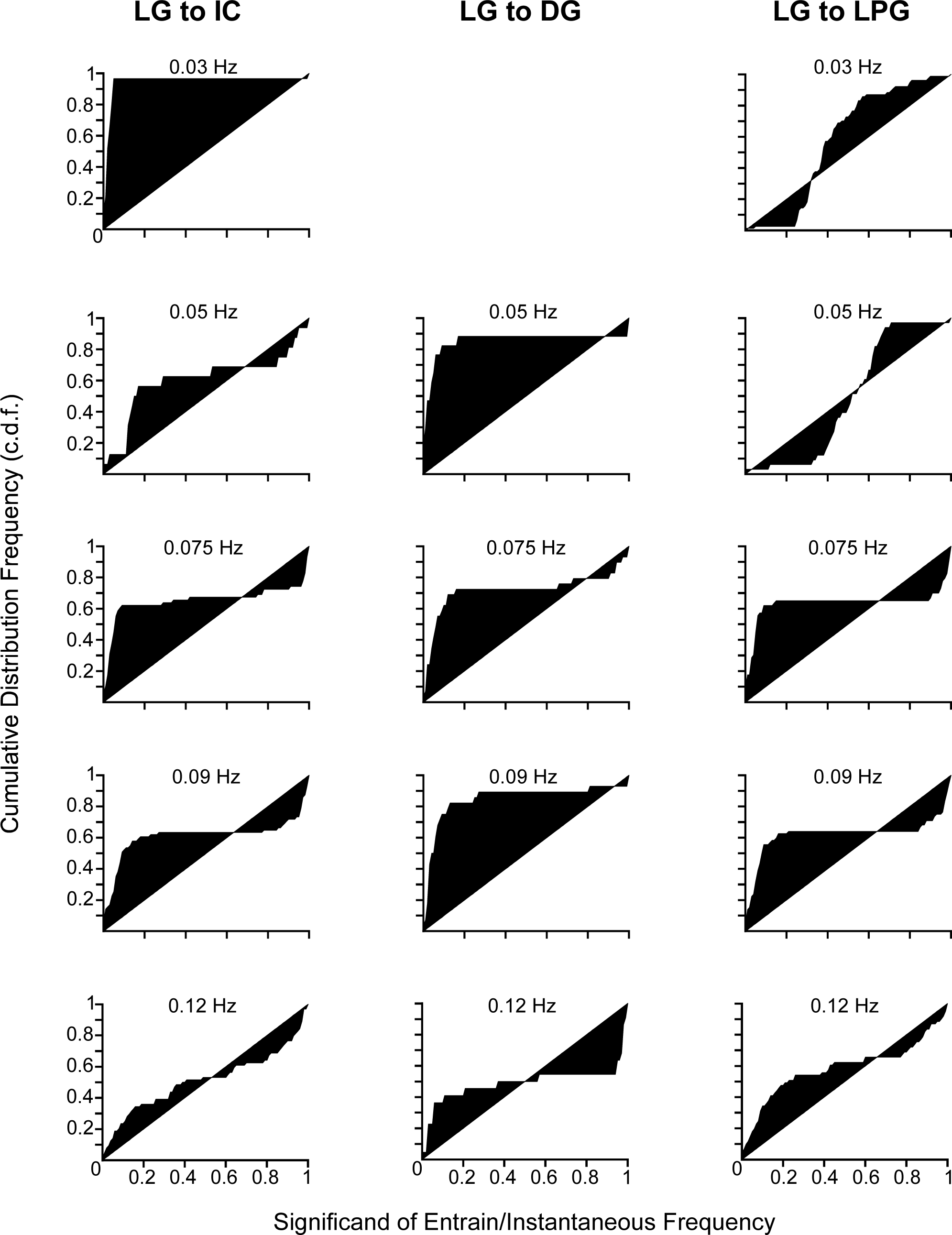
LG entrainment of IC (*left*), DG (*middle*), and LPG (*right*) during Gly^1^-SIFamide. C.d.f. graphs for IC (n = 3-7), DG (n = 2-7), and LPG (n = 2-8) neurons during LG entrainment, with each row indicating the entrainment frequency. Missing graphs indicate excluded data that was n = 1, or data that was not collected for that frequency.

**Supplemental Figure S2.**
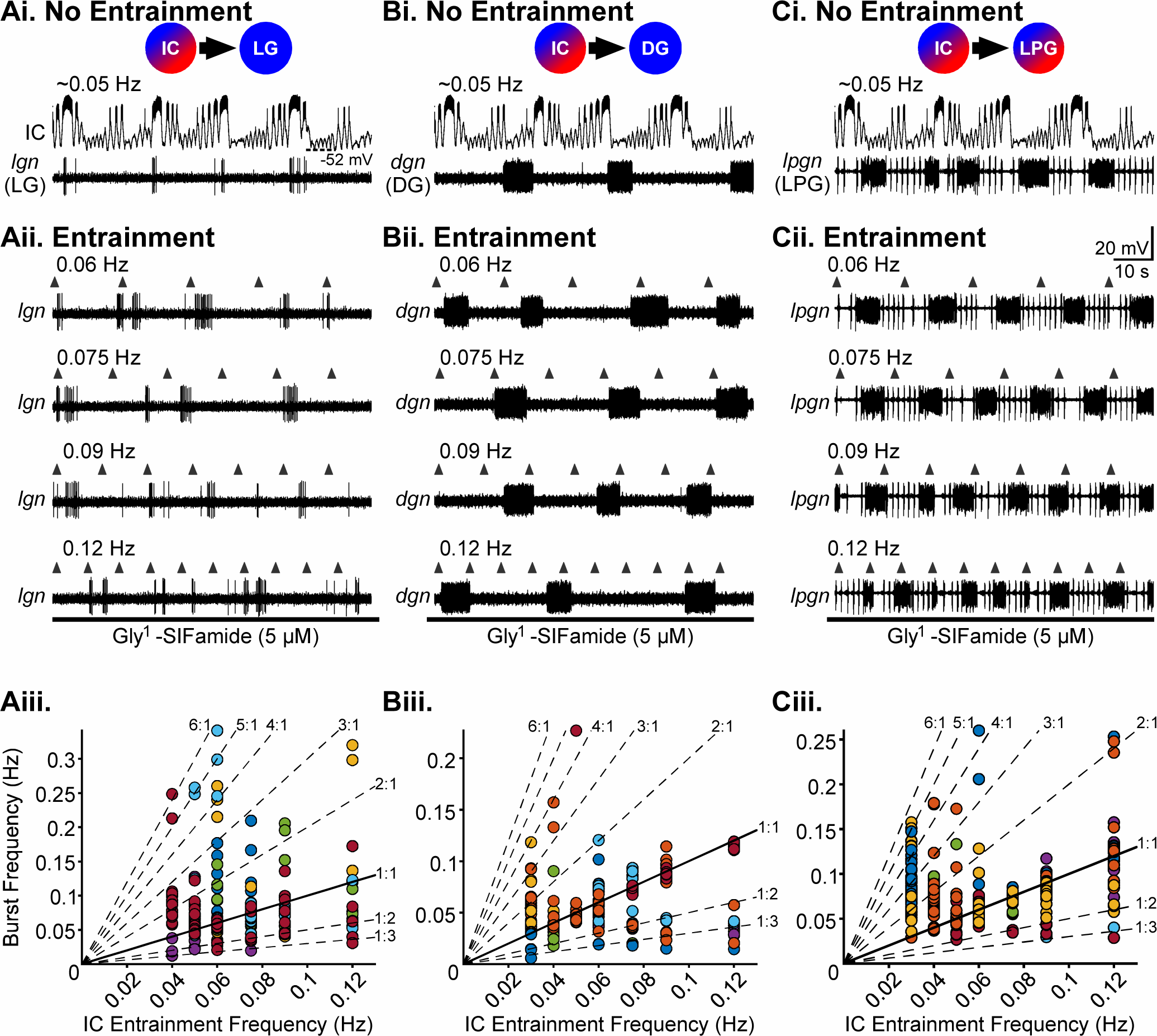
IC entrainment of LG, DG, and LPG gastric mill neurons during Gly^1^-SIFamide application. ***Ai-Ci****, **Aii-Cii**,* All columns are from the same preparation, in which IC entrainment of test neurons LG (*lgn, **Ai-iii***), DG (*dgn, **Bi-iii***), and LPG (*lpgn, **Ci-iii***) was examined. In Gly^1^-SIFamide (5 µM) prior to any manipulations, IC burst frequency was approximately 0.05 Hz and IC, DG, and LPG were bursting (***Ai-Ci***, No Entrainment). ***Aii-Cii***, To determine the ability of IC to entrain the other neurons, IC was rhythmically depolarized (*upward arrowheads*, 5 s duration; IC recordings are omitted for clarity) at different entrainment frequencies, and LG (*lgn, **Aii***), DG (*dgn, **Bii***), and LPG (*lpgn, **Cii***) activity recorded. ***Aiii-Ciii***, Instantaneous burst frequency of LG (***Aiii***), DG (***Biii***), and LPG (***Ciii***) was plotted as a function of IC entrainment frequency. Each colored data point is one instantaneous burst frequency from one preparation. Solid diagonal line indicates perfect entrainment (1:1), where instantaneous burst frequency was equal to entrainment frequency. Dashed diagonal lines indicate other modes of entrainment at which test neuron burst frequency would be a multiple, or a fraction, of entrainment frequency. Data points that fall between diagonal lines indicate a lack of IC entrainment of that neuron burst.

**Supplemental Figure S3.**
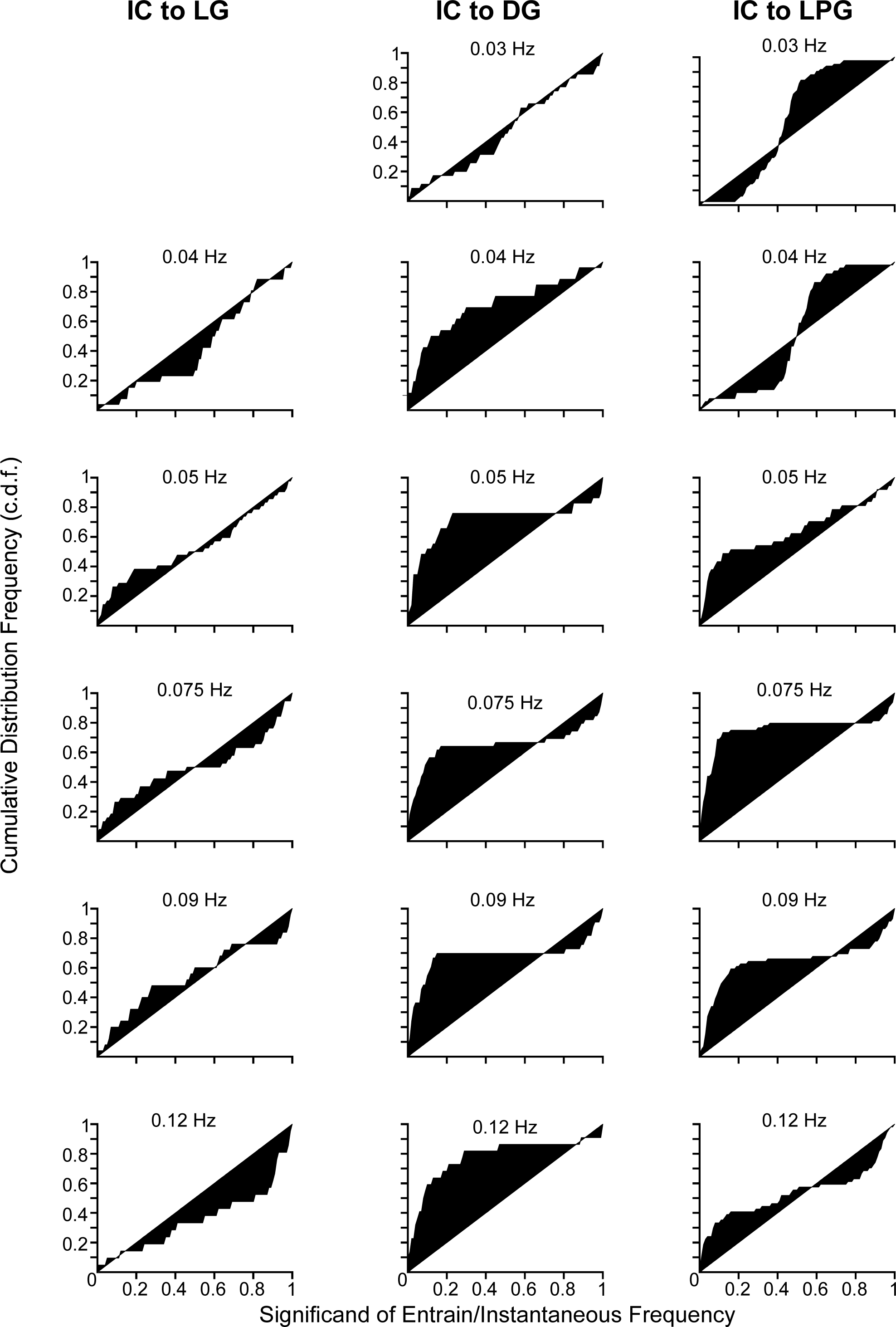
IC entrainment of LG (*left*), DG (*middle*), and LPG (*right*) during Gly^1^-SIFamide. C.d.f. graphs for LG (n = 2-6), DG (n = 3-8), and LPG (n = 2-9) neurons during IC entrainment, with each row indicating the entrainment frequency. Missing graphs indicate excluded data that was n = 1, or data that was not collected for that frequency.

**Supplemental Figure S4.**
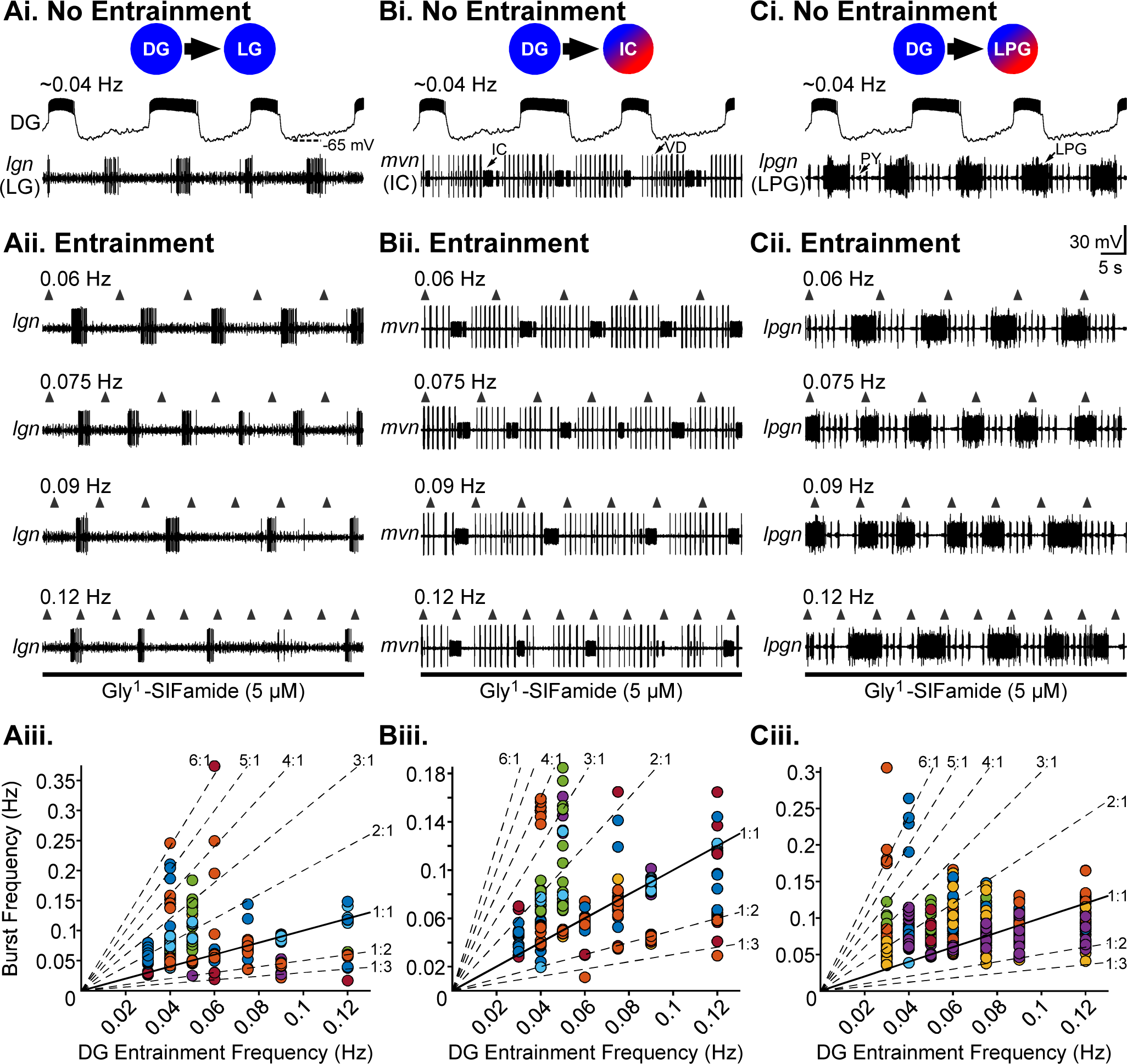
DG entrainment of LG, IC, and LPG gastric mill neurons during Gly^1^-SIFamide application. ***Ai-Ci****, **Aii-Cii***, All columns are from the same preparation, in which DG entrainment of test neurons LG (*lgn, **Ai-iii***), IC (*mvn, **Bi-iii***), and LPG (*lpgn, **Ci-iii***) was examined. In Gly^1^-SIFamide (5 µM) prior to any manipulations, DG burst frequency was approximately 0.04 Hz and LG, IC, and LPG were bursting (***Ai-Ci***, No Entrainment). ***Aii-Cii***, To determine the ability of DG to entrain the other neurons, DG was rhythmically depolarized (*upward arrowheads*, 5 s duration; DG recordings are omitted for clarity) at different entrainment frequencies, and LG (*lgn, **Aii***), IC (*mvn, **Bii***), and LPG (*lpgn, **Cii***) activity recorded. ***Aiii-Ciii***. Instantaneous burst frequency of LG (***Aiii***), IC (***Biii***), and LPG (***Ciii***) was plotted as a function of DG entrainment frequency. Each colored data point is one instantaneous burst frequency from one preparation. Solid diagonal line indicates perfect entrainment (1:1), where instantaneous burst frequency was equal to entrainment frequency. Dashed diagonal lines indicate other modes of entrainment at which test neuron burst frequency would be a multiple, or a fraction, of entrainment frequency. Data points that fall between diagonal lines indicate a lack of DG entrainment of that neuron burst.

**Supplemental Figure S5.**
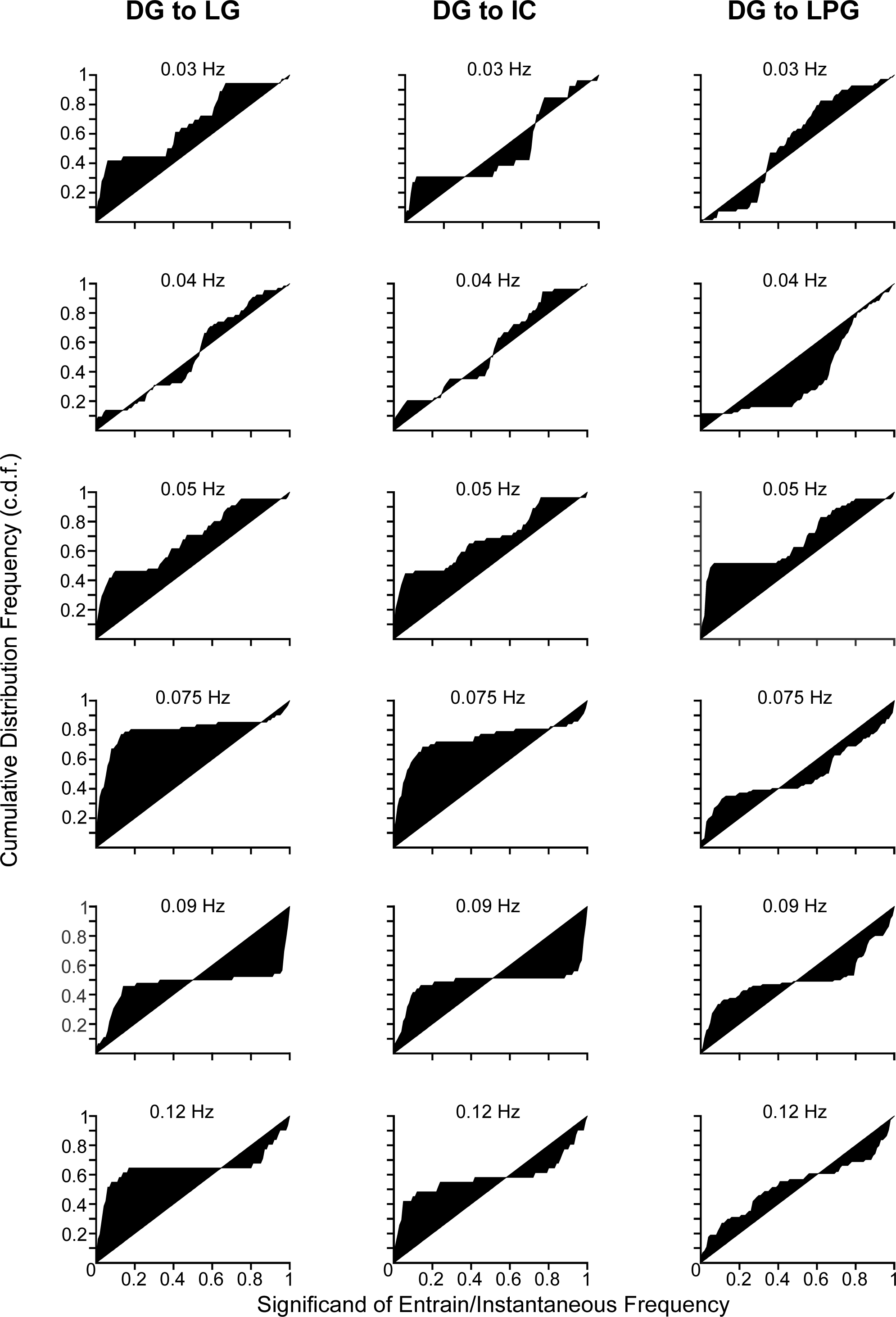
DG entrainment of LG (*left*), IC (*middle*), and LPG (*right*) during Gly^1^-SIFamide. C.d.f. graphs for LG (n = 3-8), IC (n = 2-8), and LPG (n = 3-11) neurons during DG entrainment, with each row indicating the entrainment frequency.

**Supplemental Figure S6.**
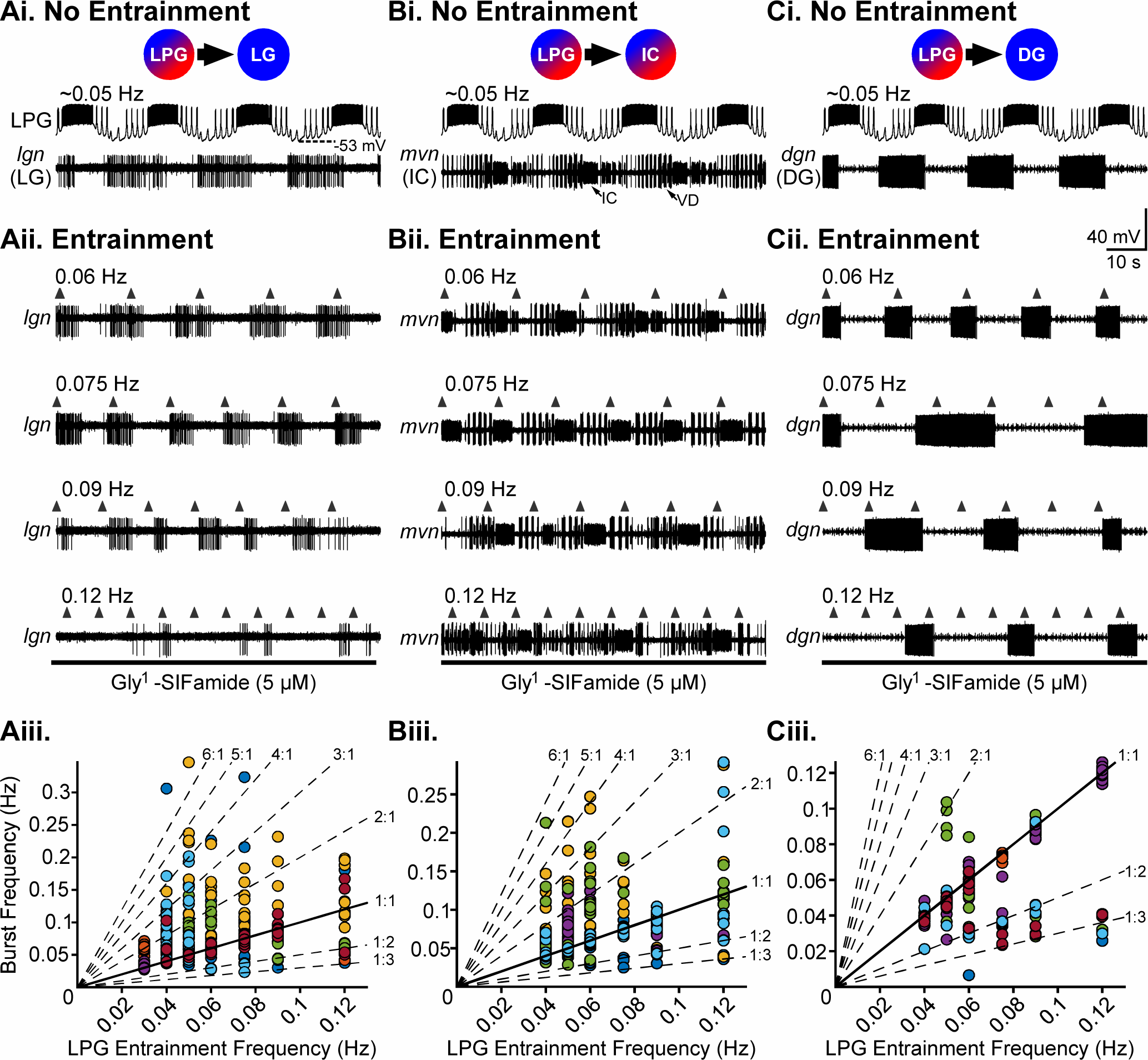
LPG entrainment of LG, IC, and DG gastric mill neurons during Gly^1^-SIFamide application. ***Ai-Ci****, **Aii-Cii***, All columns are from the same preparation, in which LPG entrainment of test neurons LG (*lgn, **Ai-iii***), IC (*mvn, **Bi-iii***), and DG (*dgn, **Ci-iii***) was examined. In Gly^1^-SIFamide (5 µM) prior to any manipulations, LPG burst frequency was approximately 0.05 Hz and LG, IC, and DG were bursting (***Ai-Ci***, No Entrainment). ***Aii-Cii***, To determine the ability of LPG to entrain the other neurons, LPG was rhythmically depolarized (*upward arrowheads*, 5 s duration; LPG recordings are omitted for clarity) at different entrainment frequencies, and LG (*lgn, **Aii***), IC (*mvn, **Bii***), and DG (*dgn, **Cii***) activity recorded. ***Aiii-Ciii***. Instantaneous burst frequency of LG (***Aiii***), IC (***Biii***), and DG (***Ciii***) was plotted as a function of LPG entrainment frequency. Each colored data point is one instantaneous burst frequency from one preparation. Solid diagonal line indicates perfect entrainment (1:1), where instantaneous burst frequency was equal to entrainment frequency. Dashed diagonal lines indicate other modes of entrainment at which test neuron burst frequency would be a multiple, or a fraction, of entrainment frequency. Data points that fall between diagonal lines indicate a lack of LPG entrainment of that neuron burst.

**Supplemental Figure S7.**
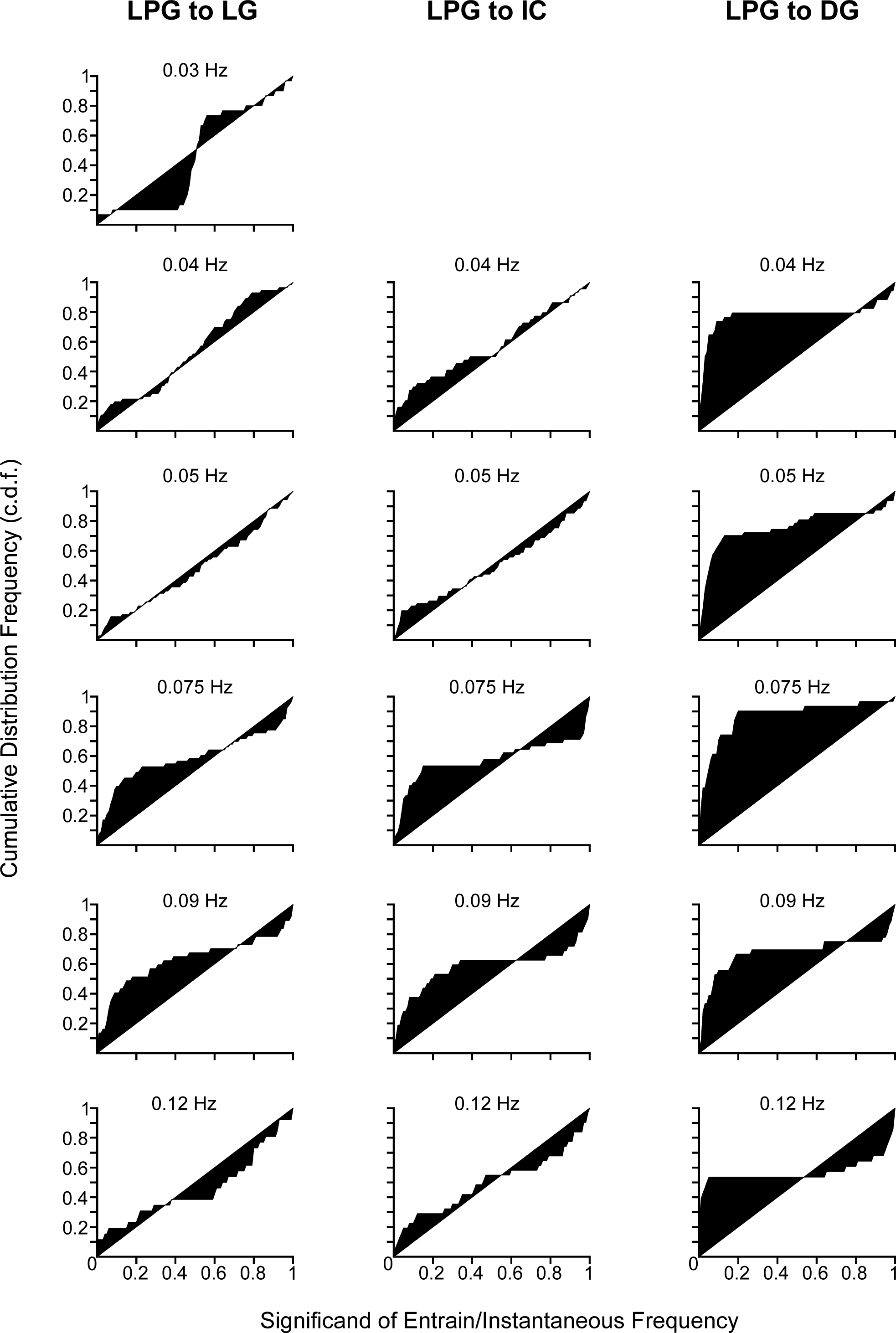
LPG entrainment of LG (*left*), IC (*middle*), and DG (*right*) during Gly^1^-SIFamide. C.d.f. graphs for LG (n = 2-6), IC (n = 4-5), and DG (n = 4-7) neurons during LPG entrainment, with each row indicating the entrainment frequency. Missing graphs indicate excluded data that was n = 1, or data that was not collected for that frequency.

**Supplemental Figure S8.**
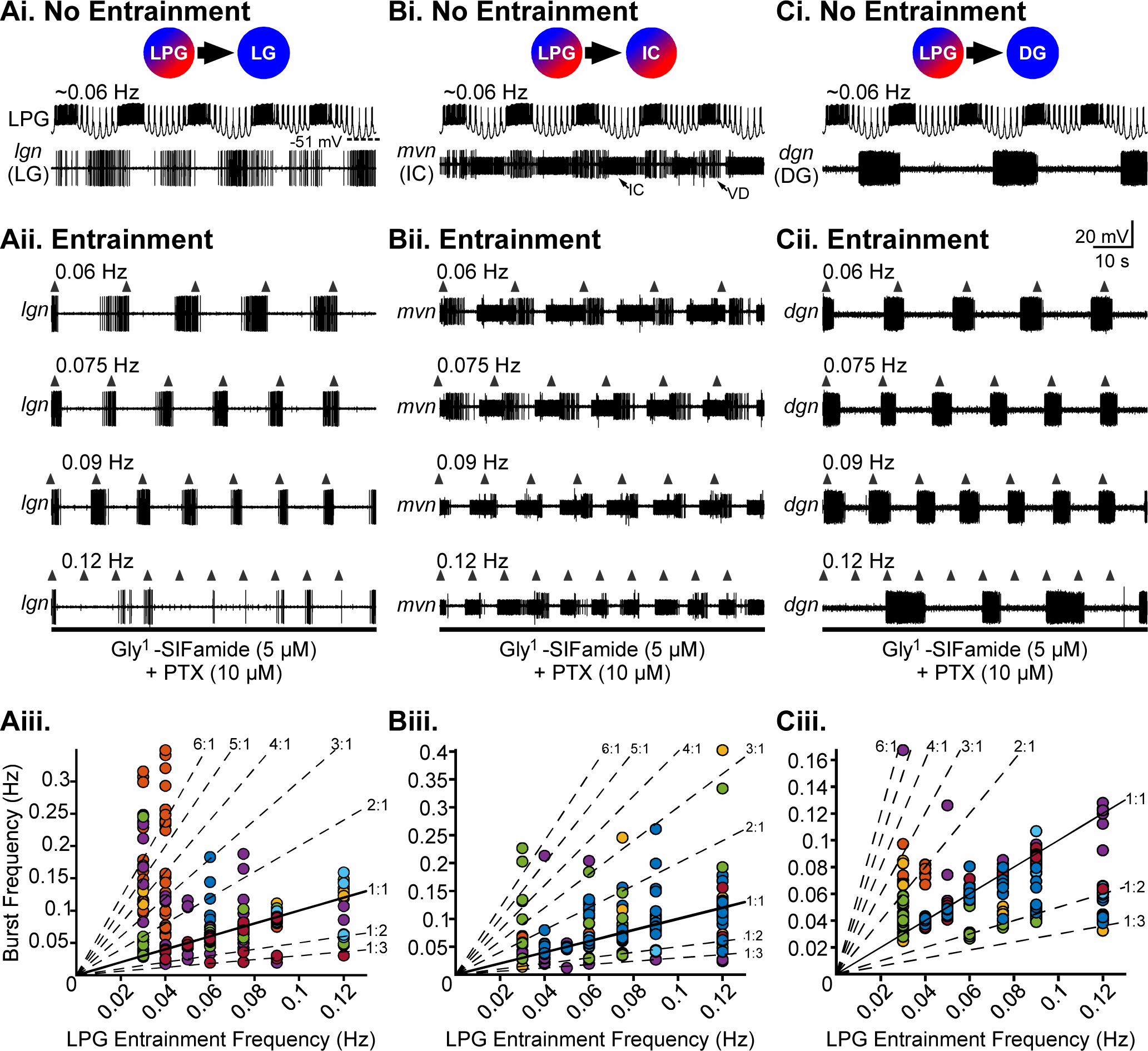
LPG entrainment of LG, IC, and DG gastric mill neurons during Gly^1^-SIFamide plus picrotoxin (SIF:PTX) application. ***Ai-Ci****, **Aii-Cii***, All columns are from the same preparation, in which LPG entrainment of test neurons LG (*lgn, **Ai-iii***), IC (*mvn, **Bi-iii***), and DG (*dgn, **Ci-iii***) was examined during Gly^1^-SIFamide (5 µM) plus picrotoxin (10 µM) (SIF:PTX). Prior to any manipulations, LPG burst frequency was approximately 0.06 Hz and LG, IC, and DG were bursting (***Ai-Ci***, No Entrainment). ***Aii-Cii***, To determine the ability of LPG to entrain the other neurons, LPG was rhythmically depolarized (*upward arrowheads*, 5 s duration; LPG recordings are omitted for clarity) at different entrainment frequencies, and LG (*lgn, **Aii***), IC (*mvn, **Bii***), and DG (*dgn, **Cii***) activity recorded. ***Aiii-Ciii***. Instantaneous burst frequency of LG (***Aiii***), IC (***Biii***), and DG (***Ciii***) was plotted as a function of LPG entrainment frequency. Each colored data point is one instantaneous burst frequency from one preparation. Solid diagonal line indicates perfect entrainment (1:1), where instantaneous burst frequency was equal to entrainment frequency. Dashed diagonal lines indicate other modes of entrainment at which test neuron burst frequency would be a multiple, or a fraction, of entrainment frequency. Data points that fall between diagonal lines indicate a lack of LPG entrainment of that neuron burst.

**Supplemental Figure S9.**
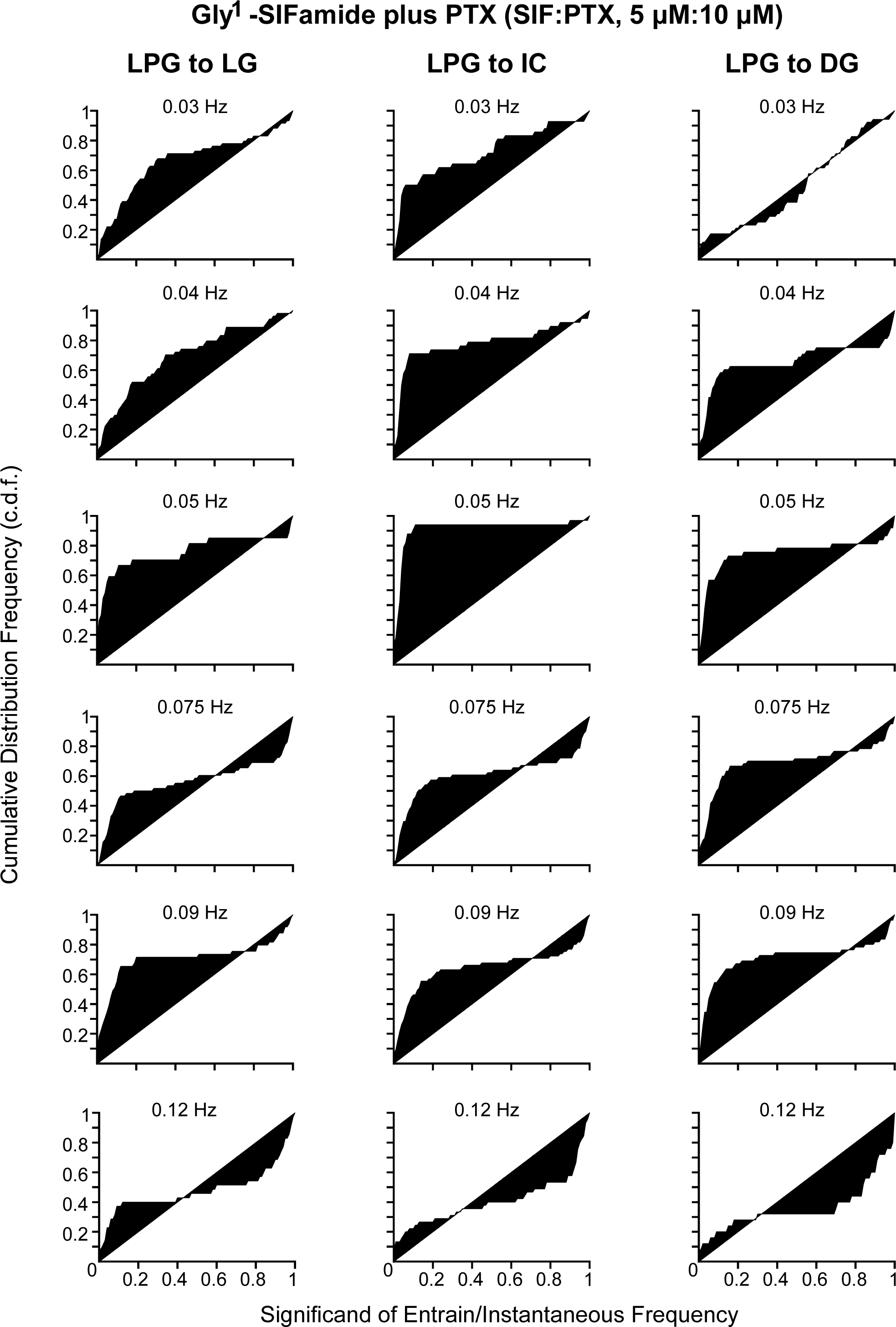
LPG entrainment of LG (*left*), IC (*middle*), and DG (*right*) during Gly^1^-SIFamide plus picrotoxin (SIF:PTX). C.d.f. graphs for LG (n = 4-7), IC (n = 4-8), and DG (n = 4-8) neurons during LPG entrainment, with each row indicating the entrainment frequency.

